# Local Glutamate-Mediated Dendritic Plateau Potentials Change the State of the Cortical Pyramidal Neuron

**DOI:** 10.1101/828582

**Authors:** Peng P. Gao, Joseph. W. Graham, Wen-Liang Zhou, Jinyoung Jang, Sergio Angulo, Salvador Dura-Bernal, Michael Hines, William W. Lytton, Srdjan D. Antic

**Author notes:** Corresponding author: Srdjan D. Antic, UConn Health School of Medicine, 263 Farmington Ave., Farmington, CT, 06030. Author Contributions: M.H., W.W.L., and S.D.A. designed the research; P.P.G., J.W.G., W.L.Z., J.J., S.A., and S.D.B. designed and performed experiments and simulations; P.P.G., J.W.G., and S.D.A. analyzed the data. The paper was written by P.P.G., M.H., W.W.L., and S.D.A., with input from all of the authors.

## Abstract

Dendritic spikes in thin dendritic branches (basal and oblique dendrites) of pyramidal neurons are traditionally inferred from spikelets measured in the cell body. Here, we used laser-spot voltage-sensitive dye imaging in cortical pyramidal neurons (rat brain slices) to investigate the voltage waveforms of dendritic potentials occurring in response to spatially-restricted glutamatergic inputs. Local dendritic potentials lasted 200–500 ms and propagated to the cell body where they caused sustained 10-20 mV depolarizations. Plateau potentials propagating from dendrite to soma, and action potentials propagating from soma to dendrite, created complex voltage waveforms in the middle of the thin basal dendrite, comprised of local sodium spikelets, local plateau potentials, and back-propagating action potentials, superimposed on each other. Our model replicated these experimental observations and made predictions, which were tested in experiments. Dendritic plateau potentials occurring in basal and oblique branches put pyramidal neurons into an activated neuronal state (“prepared state”), characterized by depolarized membrane potential and faster membrane responses. The prepared state provides a time window of 200-500 ms during which cortical neurons are particularly excitable and capable of following afferent inputs. At the network level, this predicts that sets of cells with simultaneous plateaus would provide cellular substrate for the formation of functional neuronal ensembles.

**New & Noteworthy:** In cortical pyramidal neurons, we recorded glutamate-mediated dendritic plateau potentials using voltage imaging, and created a computer model that recreated experimental measures from dendrite and cell body. Our model made new predictions, which were then tested in experiments. Plateau potentials profoundly change neuronal state -- a plateau potential triggered in one basal dendrite depolarizes the soma and shortens membrane time constant, making the cell more susceptible to firing triggered by other afferent inputs.

## Introduction

The individual spiny neuron is subjected to a variety of excitatory input patterns, often resulting from concurrent release across multiple synapses on a single dendrite. Due to combinations of temporal and spatial clustering (Kleindienst et al., 2011; Makino and Malinow, 2011; Lee et al., 2016), the amount of glutamate released on a single basilar dendrite can be quite large, potentially spilling over to extrasynaptic NMDA receptors, and temporarily overwhelming the ability of astrocytes to fully compensate (Chalifoux and Carter, 2011; Oikonomou et al., 2012). *In vitro* focal applications of comparable amounts of glutamate, or repetitive synaptic stimulations, will trigger dendritic plateau potentials in dendrites of CNS spiny neurons (Milojkovic et al., 2004; Milojkovic et al., 2005a; Major et al., 2008; Suzuki et al., 2008; Takahashi and Magee, 2009; Plotkin et al., 2011; Augustinaite et al., 2014). Originally demonstrated *in vitro*, dendritic plateau potentials have now been described *in vivo* as well (Lavzin et al., 2012; Xu et al., 2012; Smith et al., 2013; Gambino et al., 2014; Cichon and Gan, 2015; Du et al., 2017; Ranganathan et al., 2018). Glutamate-mediated dendritic spikes underlie synaptic plasticity, sensory processing, and behavior (Gordon et al., 2006; Evans et al., 2012; Lavzin et al., 2012; Cichon and Gan, 2015). Studying the biophysical aspects of dendritic plateau potentials entails experimental measurements of precise dendritic voltage waveforms along basal, oblique and tuft dendrites. These cannot be easily obtained using patch electrodes, since it is not yet possible to consistently get one or multiple electrodes onto a single thin branch (Nevian et al., 2007; Larkum et al., 2009).

NMDA spikes have been recorded by a number of groups and have been replicated computationally (Rhodes, 2006; Jadi et al., 2014; Bono and Clopath, 2017; Doron et al., 2017). By contrast, glutamate-mediated plateaus are less fully explored, and less well understood. In this paper, we analyze properties and mechanisms of dendritic plateaus through coupled experimentation and computer simulation. New experimental measurements in dendrites of cortical pyramidal neurons were used to create a detailed computational model of dendritic plateaus in a full morphology model of a Layer 5 cortical pyramidal cell, demonstrating that the dynamics of these potentials will have a major effect on spike generation: the plateau not only places the cell closer to spike threshold but also shortens the membrane time constant. As a consequence, other incoming excitatory postsynaptic potentials (EPSPs), arriving on other dendritic branches, become much more potent drivers of action potential (AP) initiation during the plateau event. The results from the detailed model support a theoretical framework in which plateau potentials allow cortical pyramidal neurons to respond quickly to ongoing network activity and potentially enable synchronized firing to form active neural ensembles (Legenstein and Maass, 2011; Chiovini et al., 2014; Antic et al., 2018).

## Methods and Materials

### Brain slice and electrophysiology

Sprague Dawley rats (P21 – 42) of both sexes were anesthetized with isoflurane, decapitated, and the brains were removed with the head immersed in ice-cold, artificial cerebrospinal fluid (ACSF), according to an institutionally approved animal protocol. ACSF contained (in mM) 125 NaCl, 26 NaHCO_3_, 10 glucose, 2.3 KCl, 1.26 KH_2_PO_4_, 2 CaCl_2_ and 1 MgSO_4_, at pH 7.4. Coronal slices (300 μm) were cut from frontal lobes. Whole-cell recordings were made from visually identified layer 5 pyramidal neurons. Intracellular solution contained (in mM) 135 K-gluconate, 2 MgCl_2_, 3 Na_2_-ATP, 10 Na_2_-phosphocreatine, 0.3 Na_2_-GTP and 10 Hepes (pH 7.3, adjusted with KOH). In some experiments, the intracellular solution was enriched with the fluorescent dye Alexa Fluor 594 (40 μM) to aid positioning of glutamate stimulation electrodes on distal dendritic branches. Electrical signals were amplified with a Multiclamp 700A and digitized with two input boards: (1) Digidata Series 1322A (Molecular Devices, Union City, CA) at 5 kHz, and (2) Neuroplex (RedShirtImaging, Decatur, GA) at 2.7 kHz sampling rate. All experiments were performed at 33-34°C. Glutamate microiontophoresis was performed using sharp pipettes (40 ± 10 MΩ) pulled from borosilicate glass with filament (1.5 mm OD), and backfilled with 200 mM Na-glutamate (pH=9). A programmable stimulator (Clampex, Molecular Probes) and stimulus isolation unit IsoFlex (A.M.P.I.) were used to iontophoretically eject glutamate. A motorized micromanipulator Shutter M-285 was used to drive the tips of glutamate pipettes into the slice tissue with both “X” and “Z” axis motors engaged simultaneously. This was done to prevent bending of the sharp electrode, which occurs if a simple “Z” axis motion is used.

### Dye Injections

The voltage-sensitive dye injection protocol was previously described in ref. (Antic, 2003). Briefly, neurons were filled through whole-cell recording pipettes with a styryl voltage-sensitive dye JPW3028 (Potentiometric Probes, Farmington, CT) dissolved in standard K-gluconate based intracellular solution. Dye loading patch pipettes were filled with two varieties of the same intracellular solution; one with and one without the dye. Dye-free solution occupied the very tip of the pipette, while the back of the pipette lumen was microloaded with dye-rich solution (400 – 800 μM). The purpose of dye-free solution in the tip of the patch pipette was to prevent dye-leak during the maneuver through brain slice tissue. JPW3028 is lipophilic and binds indiscriminately and irreversibly to all membranes. Even a small amount of dye leak during the formation of the gigaohm seal can generate strong fluorescent background, which has a devastating effect on dendritic optical signals. The filling pipette was carefully pulled out (outside-out patch) and brain slices were left to incubate for 40-120 minutes at room temperature. Just before optical recordings, the cells were re-patched with dye-free pipette at physiological temperature (33-34^°^C).

### Dendritic voltage imaging

Voltage-sensitive dye imaging was performed on a Zeiss Axioskop 2FS microscope equipped with NeuroCCD camera (RedShirtImaging). We used Zeiss 40X objective IR-Achroplan 0.80 NA. Laser spot illumination was used to excite the voltage-sensitive dye (Zhou et al., 2007). Into the epi-illumination port of the microscope, we inserted a F200 μm fiber optic guide with a collimator. The laser beam (Cobolt Samba 532 nm, 150 mW) was focused on the other side of the fiber optic guide using a microscope objective lens. This arrangement produced a motionless spot of laser light (25 - 50 μm in diameter) at the object plane. A region of interest (ROI) was brought into the laser spot using X-Y microscope platform. The laser beam was interrupted by an electro-programmable shutter (Uniblitz, Vincent Associates). Laser beams were directed onto the preparation with the help of Zeiss epi-illumination filter cube: exciter 520 ± 45 nm; dichroic 570 nm; emission >610 nm.

Optical signals were recorded with 80 x 80 pixels at a 2.7 kHz frame rate, stored, and then temporally filtered (off-line) with digital Gaussian low-pass filter (1050 Hz cut-off), and Butterworth high-pass filter (4.5 Hz), unless otherwise specified. To improve signal-to-noise ratio several pixels (3 – 8 pixels) were selected inside the region of interest and spatially averaged, unless otherwise specified. With the 40X magnification lens, used in this study, each pixel covers 4.8 x 4.8 μm in the object field. After the experiment, fluorescent images were captured with IR-1000 Dage CCD camera. In order to obtain whole-field photographs of the dendritic tree, brain slices were removed from the recording chamber and mounted on a microscope slide in water-based mounting medium. Mounted microscope slides were transferred to Zeiss Axiovert 200M imaging station where photographs were taken with AxioVision LE system using 20x dry and 40x oil immersion objectives.

### Physiology data analysis

Optical and electrical measurements were analyzed using the software Neuroplex (RedShirtImaging) and Clampfit (Molecular Probes). Plateau amplitude was measured as a difference between the peak depolarization after the last AP in the burst and the baseline. Duration of the plateau depolarization was measured at 50% of plateau amplitude. The linear correlation coefficient (c.c.) and graph plotting were done in custom made software written in Python. The backpropagation action potential (bAP) amplitude atop plateaus was measured from the plateau phase (‘p’) to the peak of APs. The amplitude of the test-pulse-evoked voltage transients (ΔVm) was measured as a difference between voltage transient peak and baseline established just prior to current injection (plateau level). Membrane time constant (TAU) was measured in Clampfit by fitting an exponential through the charging curve.

### Modeling

The simulations were implemented with the NEURON simulator (version 7.5) (Hines and Carnevale, 1997) through its Python interface (version 2.7). The full model is available from ModelDB (accession number 249705). Here briefly, the multi-compartment cell was modified from a morphologically detailed L5 pyramidal neuron (Acker and Antic, 2009). It has 85 compartments and 439 segments in total: 36 compartments for basal dendrites, 45 for apical dendrites, 3 for soma and 1 for axon.

The detailed channel parameters are listed in Table 1.

**Table 1.** Model parameters.

| Table 1 |  |  |  |
| --- | --- | --- | --- |
| Channels | Conductance | Units | Ref |
| Voltage-gated sodium channels | | pS/$\mu\text{m}^2$ | (Acker and Antic, 2009)<br>(Mainen et al., 1996) |
| - Soma | 900 |  |  |
| - Basal dendrite | $150 - \text{distance} * 0.5$ | | |
| - Apical dendrite | 375 |  |  |
| - Axon | 150 |  |  |
| - AIS | 5,000 |  |  |
| A-type potassium channels | | pS/$\mu\text{m}^2$ | (Migliore et al., 1999; Acker and Antic, 2009) |
| - Soma | 150 |  |  |
| - Basal dendrite | $\text{Distal } (150 + 0.7 * \text{distance}) * (1/300 * \text{distance})$ | | |
|  | Proximal (150 + 0.7*distance) * (1 - 1/300*distance) |  |  |
| - Apical dendrite | Distal 300* (1/300*distance)<br>Proximal 300* (1 - 1/300*distance) |  |  |
| High voltage activated calcium channels | | pS/$\mu\text{m}^2$ | (Mainen et al., 1996; Kampa and Stuart, 2006; Acker and Antic, 2009) |
| - Soma | 2 |  |  |
| - Basal dendrite | Distal 0.4<br>Proximal 2 |  |  |
| - Apical dendrite | Distal 0.4<br>Proximal 2 |  |  |
| Low voltage activated calcium channels | | pS/$\mu\text{m}^2$ | (Mainen et al., 1996; Kampa and Stuart, 2006; Acker and Antic, 2009) |
| - Soma | 2 |  |  |
| - Basal dendrite | Distal 1.6<br>Proximal 2 |  |  |
| - Apical dendrite | Distal 1.6<br>Proximal 2 |  |  |
| HCN channels |  | S/cm<sup>2</sup> | (Kole et al., 2006) |
| - Soma | 0.0001 |  |  |
| - Basal dendrite | 0.0001 |  |  |
| - Apical dendrite | $0.0002 * (-0.8696 + 2.0870 * \exp(\text{distance}/323))$ | | |
| Calcium activated potassium channels | 2.68e-4 | S/cm<sup>2</sup> | (Reetz et al., 2014; Stadler et al., 2014) |
| Kv channel | | pS/$\mu\text{m}^2$ | (Mainen et al., 1996; Acker and Antic, 2009) |
| - Soma | 40 |  |  |
| - Basal dendrite | 40 |  |  |
| - Apical dendrite | 40 |  |  |
| - Axon | 100 |  |  |

### Modeling bAPs

We investigated the properties of bAPs on 6 different basal branches using similar methods as (Acker and Antic, 2009). Square waveform current pulses (3 nA, 1.75 ms) were injected in the soma; and the spike amplitude and peak time were measured at different locations along 6 basal dendrites. For TTX and 4-AP conditions, the sodium channel or A-type potassium channel conductances were set to 0, respectively. The binned average (bin size = 20 μm) spike amplitude and peak latency were plotted against the location on basal dendrites, as the distance from soma.

### Modeling glutamate inputs

The glutamate microiontophoresis experiments were simulated by activation of AMPA and NMDA receptor models. AMPA and NMDA receptors were activated within a dendritic segment (∼60 μm length) on the targeted basal dendrite. The proximal edge of the activated dendritic segment was 70 μm away from the soma. In order to study the effect of input location and the spatial profile in dendrites, the length of the active dendritic segments was decreased to 10 - 30 μm.

Glutamate receptor channels were divided into two groups: synaptic (AMPA and NMDA, ratio 1:1); and extrasynaptic (NMDA). The ratio of synaptic to extrasynaptic NMDA conductance was set at 1:1. The extrasynaptic NMDA receptors were always activated 5 ms after the activation of the neighboring synaptic NMDA receptors. The glutamate input strength was regulated through a “weight factor”, which simultaneously scaled three parameters: [1] the number of activated receptors; [2] the synaptic weights; and [3] the receptor activation time window. In our conceptual model, clustered synaptic inputs overcome the capacity of the glutamate uptake, thus causing glutamate molecules to linger longer in the vicinity of the receptors. For this reason, in the current model (which nicely matches several sets of experimental data, see below) an increase in glutamate input intensity causes glutamate receptors to spend longer time in activated state. The minimum receptor activation time window was 40 ms when the weight factor was set to 0, and the maximum time window was 90 ms, when the weight factor was equal to 1. Two types of temporal activation patterns were used: [1] the random function (continuous uniform distribution) and [2] the beta random function, available in the Python NumPy library. Two membrane mechanisms for NMDAr were used, resulting in two cell models: [Model 1] the classical two state kinetic model with alpha – function (Destexhe et al., 1994); and [Model 2] the triple-exponential “envelope” time course and a sigmoidal voltage dependence (Major et al., 2008). The same AMPA receptors were used in Models 1 & 2 (Destexhe et al., 1994). In all simulations, the maximum conductances (g_max) for AMPA, synaptic NMDA and extrasynaptic NMDA were 0.05 μS, 0.005 μS, and 0.005 μS, respectively. Barrages of EPSPs were simulated as the activation of AMPA receptors only on 5 dendritic branches, different from the dendritic branch receiving glutamate input for initiation of local plateau potential.

### Quantifications of the modeling results

The measurements of plateau amplitude, duration and spikes per plateau in both experiments and simulations were implemented using Python. Plateau amplitude was calculated as the minimum voltage value between the last two APs riding on top of the plateau potential. Plateau duration was calculated as the period of time during which the voltage was higher than the half plateau amplitude (half-width). The membrane time constant was measured in response to a square wave current injection, as the time required for decaying to 37% of its initial value. Local input resistance in dendrites was attained by injecting a standard current pulse (100 pA, 100 ms) and measuring local ΔVm.

## Results

We simultaneously recorded dendritic and somatic potentials in 10 layer V pyramidal neurons, with somatic recordings alone in an additional 15 cells. Simulation fitting was done by hand in a single full multicompartment morphology over several thousand simulation runs.

### I Experiment: glutamate-evoked dendritic plateau potentials

Plateau potentials were induced by brief (5 ms) pulses of iontophoretically ejected glutamate applied locally on individual dendrites of cortical Layer 5 pyramidal neurons (Fig. 1*A1, Camera 1*, glut.). Voltage waveforms of plateau potentials in basal and oblique dendrites were recorded with voltage-sensitive dye imaging (Fig. 1*A2*, dend). Dendrites were illuminated by a spot of laser light, and the image of a dendritic segment was projected onto a fast camera (Fig. 1*A1, Camera 2*). Simultaneously with dendritic voltage imaging, we recorded somatic membrane potential in whole-cell (Fig. 1*A2*, soma).

**Fig. 1.**
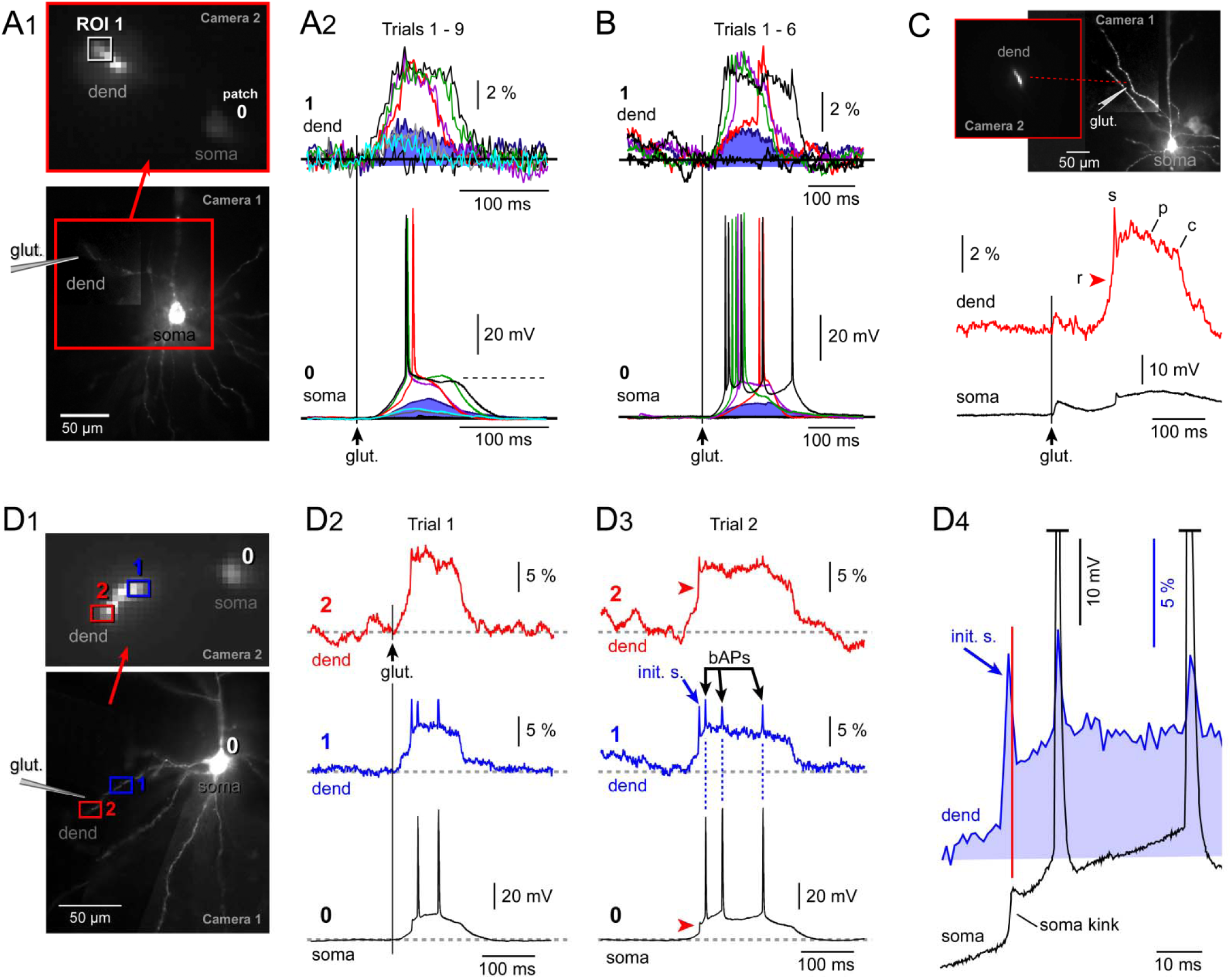
Dendritic voltage waveforms. **Voltage waveforms of the glutamate evoked dendritic plateau potentials**. **(A**_**1**_) Bottom image: Pyramidal neuron filled with JPW-3028. Image was acquired by standard camera used for patching (Camera 1). Top image: Only one dendrite was illuminated by a laser spot illumination technique, and imaged by a low-resolution (80×80 pixel), fast (2.7 kHz), voltage-imaging camera (Camera 2). **(A**_**2**_) Glutamate was applied iontophoretically (5 ms) at location indicated in *A*_*1*_, bottom image. The intensity of the glutamatergic stimulation was increased in equal steps through 9 recording trials (Trials 1-9), while optical signals were recorded in the dendritic region of interest (ROI 1) marked by rectangle in *A*_*1*_, top image. “Soma” marks the somatic whole cell recordings obtained simultaneously with the optical signals. (**B**) Same experimental paradigm as in *A*, except different cell. (**C**) Glutamatergic stimulation of an oblique dendrite (duration 5 ms) triggers a local plateau potential. “r” - fast rise; “s” - initial sodium spikelet; “p” - plateau phase; “c” – collapse, of the somatic plateau potential. (**D**_**1**_) Pyramidal neuron filled with JPW-3028. The position of the glutamate-filled sharp electrode is marked by drawing. (**D**_**2**_) In Trial 1, glutamate-induced membrane potential changes are recorded simultaneously at two ROIs on the same dendrite and in the cell body (whole-cell). (**D**_**3**_) In Trial 2, the glutamatergic stimulus was increased by 20% causing a longer lasting plateau phase. The initial spikelet in the dendritic recording (blue arrow) did not cause and AP in somatic recording, but instead it caused a rapid inflection preceding the plateau phase (red arrowhead). Three backpropagating APs (“*bAPs”*) recorded in dendrite are marked by black arrows. (**D**_**4**_) Blowup of *D*_*2*_ on a faster time scale, to show that initial spikelet (init. s.) precedes the somatic kink.

#### Glutamate threshold

The glutamate pulse invariably produced a dendritic depolarization (Fig. 1, *A* and *B*) in all neurons tested in this way (n = 10). At lower glutamate input intensities, the depolarization was seen as a brief EPSP (Fig. 1*A2*, blue shading). As the intensity of the glutamatergic stimulation was gradually increased in equal steps (∼10 nA, intensity of iontophoretic current) a threshold was reached with a discontinuity from small EPSP depolarizations up to the plateau voltage level. The passage from subthreshold (Fig. 1, *A2* and *B*, dark blue trace) to suprathreshold (red trace), could be seen in both dendrite and soma. Further increase in glutamate input intensity did not result in greater depolarization, but prolonged the duration of the plateau (Fig. 1, *A* and *B*, trials 1 – 9), suggesting an all-or-none spike mechanism (Schiller et al., 2000).

### Voltage waveform at the input site

Locally, dendritic plateaus showed a characteristic voltage waveform (Fig. 1*C*), beginning with a rapid onset (‘r’). The sudden increase in voltage, seen clearly in the dendrite, was low-pass filtered by the dendritic cable and so appeared less abrupt in the soma (Fig. 1*C*, soma). This onset phase was capped with an initial sodium spikelet (‘s’), described further below. The dendritic plateau phase (‘p’) lasted >100 ms but terminated with an abrupt decline, or collapse (‘c’) back to resting membrane potential (RMP).

### Superposition of 3 spikes in mid dendrite

Simultaneous recordings from 2 dendritic sites (Fig. 1*D1*) revealed three varieties of dendritic spikes including: [1] square-shaped glutamate-mediated dendritic plateau potentials (Fig. 1*D2*, red trace); [2] dendrite-originating fast sodium spikes uncoupled from somatic APs (Fig. 1*D3*, “*init. s*”); and [3] fast sodium spikes associated with firing of APs in the cell body – backpropagating APs (Fig. 1*D3*, “*bAPs*”). The most distal dendritic segment, closest to the point of glutamatergic input was dominated by square-shaped plateau potentials (Fig. 1, *D2* and *D3*, red traces). The cell body was dominated by APs (black traces). The mid dendritic segment experienced a complex voltage waveform resulting from a superposition of the three aforementioned spike varieties: [1] plateau potential, [2] initial spikelet, and [3] bAPs (Fig. 1, *D2* and *D3*, blue traces). To determine direction of propagation, we examined dendritic and somatic records on a faster time scale (Fig. 1*D4*). The conclusions of this analysis are laid out in the next paragraph.

### Initial Na^+^ spikelet

The first peak in the dendritic voltage waveform was not a bAP, but rather a dendritic fast sodium spikelet at the beginning of the plateau that propagated orthodromically from dendrite to cell body. Initial spikelets (‘init. s.’) invariably failed to trigger a somatic or axonal AP, but were clearly seen as kinks in somatic recordings (Fig. 1*D4*, soma kink). The peak of the dendritic sodium spikelet (‘init. s.’) occurred prior to its appearance in the soma (red vertical line). This spikelet has been previously shown to disappear in the presence of the sodium channel blocker, TTX (Milojkovic et al., 2005b; Nevian et al., 2007).

### Backpropagating action potentials (bAPs)

Simultaneous dendritic and somatic recordings showed that plateau-triggered APs occurred in the cell body before the dendrite, thus indicating that somatic APs propagated from soma into the dendrites, riding atop the plateau potential (Fig. 1*D4*). We found that bAPs could partially invade distal dendritic segments of basal dendrites even during local plateau depolarizations (Fig. 1, *D2* and *D3*, blue and red traces). bAP voltages were diminished in amplitude at the distal site (ROI-2) compared to the proximal site (ROI-1), demonstrating the retrograde direction of propagation, from soma to dendrite. It is important to emphasize that the initial spikelet (Fig. 1*D3*, “*init. s*”) showed a completely opposite trend: initial spikelet voltages were diminished in amplitude at the cell body (ROI-0, soma) compared to the dendrite site (ROI-1, dend) due to their orthograde direction of propagation, from dendrite to soma. Fast sodium spikelets propagating from dendrite to soma encounter a strong current sink imposed by the large amount of leaky membrane contained in the cell body and basal dendrites. The dendrite-originating initial spikelets fail to charge the cell body rapidly or sufficiently, thus failing to invade the soma (Goldstein and Rall, 1974; Moore and Westerfield, 1983).

#### II Simulation: computer model constrained by experimental measures

We built a full neuron model (Fig. 2, *A1* and *A2*) with basal dendrites tuned to reproduce previously published data. More specifically, these simulations matched experimental studies of bAPs (without plateaus) obtained previously (Antic, 2003; Acker and Antic, 2009), showing a ∼18 mV/100 μm AP peak amplitude decrease with distance; AP backpropagation velocity: ∼180 μm/ms (Fig. 2, *B1* - *B3*). Elimination of voltage-gated Na^+^ channels (TTX condition) in model dendrites, increased attenuation and decreased delay, while removal of A-type K^+^ channels (4-AP condition) had opposite effects, matching bAP amplitude decrement with TTX or 4-AP, measured experimentally (Acker and Antic, 2009).

**Fig. 2.**
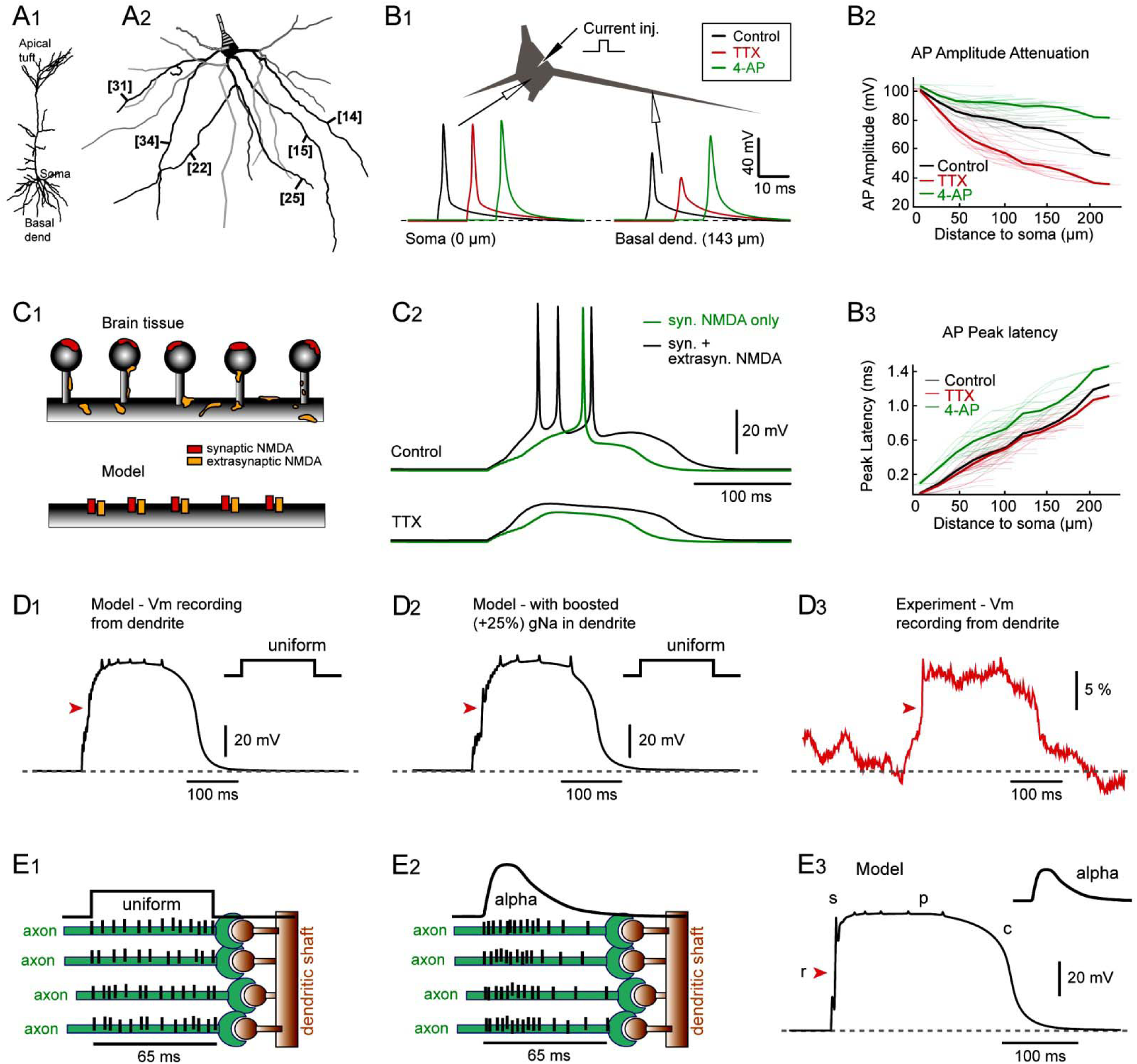
Model outline, AP backpropagation, glutamate spillover and temporal clustering of inputs. **Model outline**.**(A**_**1**_) Reconstructed pyramidal neuron. **(A**_**2**_) Six basal dendrites, mostly used in data quantifications, are labeled by bracketed numbers. **(B**_**1**_) The bAP amplitude, peak latency, response to TTX and 4-AP match the experimental measurements obtained in basal dendrites using voltage sensitive dyes. (**B**_**2**,**3**_) Quantification of the model results. **(C**_**1**_) Glutamate stimulations activate the AMPA and NMDA receptors on dendritic spines (red) and the NMDA receptors on extrasynaptic surfaces, including the spine head, spine neck, and dendritic shaft (yellow). **(C**_**2**_**) (upper)** The comparison of simulated traces with and without extrasynaptic NMDARs. Green = Somatic response evoked by synaptic NMDARs only. Black - Somatic response evoked by conjugate activation of both synaptic and extrasynaptic NMDARs. **(lower)** Same traces as in *C*_*2*_*-upper*, but with the application of TTX to block all voltage-activated sodium channels. **(D**_**1**_) Membrane potential change in basal dendrite during a plateau potential obtained in model cell shown in panel *A*. **(D**_**2**_) Same as in *D*_*1*_, except gNa_bar increased by 25%. **(D**_**3**_) Membrane potential change in basal dendrite obtained by voltage-sensitive dye imaging (*from Fig*.*1D*_*3*_). **(E**_**1**_) Temporal organization of the glutamatergic inputs – uniform random within time window of 65 ms. Presynaptic axons (green) impinging on dendritic spines (brown). Ticks indicate APs in axon. **(E**_**2**_) Temporal organization of inputs – alpha random function. **(E**_**3**_) Dendritic plateau potential with alpha distribution of inputs. “r”, “s”, “p”, & “c” same definition as in Fig. 1.

Plateau potential activation is thought to be the result of synaptic activation on spine heads, as well as glutamate-spillover activation of extrasynaptic NMDARs on spine heads and necks, and on dendritic shafts (Fig. 2*C1*) (Arnth-Jensen et al., 2002; Scimemi et al., 2004; Chalifoux and Carter, 2011; Oikonomou et al., 2012). We did not explicitly model diffusion of glutamate but instead provided a delay of 5 ms to activate these receptors at the same location on the model dendrites (Fig. 2*C1*, bottom). A plateau potential could be obtained with just synaptic NMDAR activation (Fig. 2*C2*, Control, green trace), but showed increased initial slope, amplitude, duration, and number of APs with the addition of extrasynaptic NMDAR activation (Fig. 2*C2*, Control, black trace). Plateau potential generation was dependent on adequate NMDAR activation and could be replicated in any oblique or basilar dendrite in both full morphology or in simplified neurons with one or more basilar/oblique dendrites. It was therefore robust to other changes in local ion channel densities, such as the elimination of Na^+^ channels (Fig. 2*C2*, TTX). It was also robust to moderate changes in NMDA parameters, including activation and inactivation time constants, and glutamate activation duration, and to glutamate stimulation location (see below). Simulations reproduced major behaviors seen experimentally in the present study (Fig. 1) including: (i) an inflection point on the rising phase at glutamate threshold (Fig. 2, *D1* and *D2*, red arrowheads); (ii) plateau phase duration of 200-500 ms; (iii) the initial spikelet; (iv) bAPs on plateau, and (v) abrupt plateau collapse.

Approximately 13% of basal dendrites are endowed with an ability to generate local sodium spikelets (Milojkovic et al., 2005b); most likely due to higher local concentrations of Na^+^ channels. Increasing the maximum sodium channel conductance (gNa_max) in basal dendrites by 25% had only minor effect on the plateau morphology, but improved initial spikelet (Fig. 2*D2*). Next, we considered the temporal organization of incoming glutamatergic inputs impinging on the dendrite: comparing uniform random temporal distribution of synaptic activation (Fig. 2*E1*, “uniform”) versus grouping of synaptic inputs at the beginning of temporal window (Fig. 2*E2*, “alpha”). The alpha-pattern grouping of the excitatory inputs improved the resemblance between model (Fig. 2*E3*) and experimental measurement (Fig. 2*D3*), producing a more abrupt initial rise ‘r’ (red arrowheads).

### Increased plateau duration with increased stimulation

In the next series of experiments, voltage-sensitive dye was replaced by Alexa Fluor 594, glutamate electrode positioned on basal dendrite, and recordings obtained in the cell body only. Increasing the intensity of the dendritic glutamatergic input (5 ms glutamate ejection on basal dendrites; 70–110 μm from soma center; n=15) in equal increments produced characteristic families of traces (Fig. 3A), which could be reproduced by simulation (Fig. 3B). In real neurons, the plateau amplitude at cell body (slow component of somatic depolarization) showed a sigmoidal relation to the intensity of glutamatergic input presented on the dendrite (Fig. 3*C1*, green). Similar distribution of the somatic plateau amplitudes was produced by our model (Fig. 3*C1*, red). Despite the plateau amplitude saturation (Fig. 3*C1*), cortical pyramidal neurons can interpret additional increases in glutamate input intensity by the means of plateau duration and AP firing. In both experiment and model, plateau duration increased linearly with increasing glutamate stimulation (Fig. 3*C2*), which in turn caused an increase in AP count (Fig. 3*C3*).

**Fig. 3.**
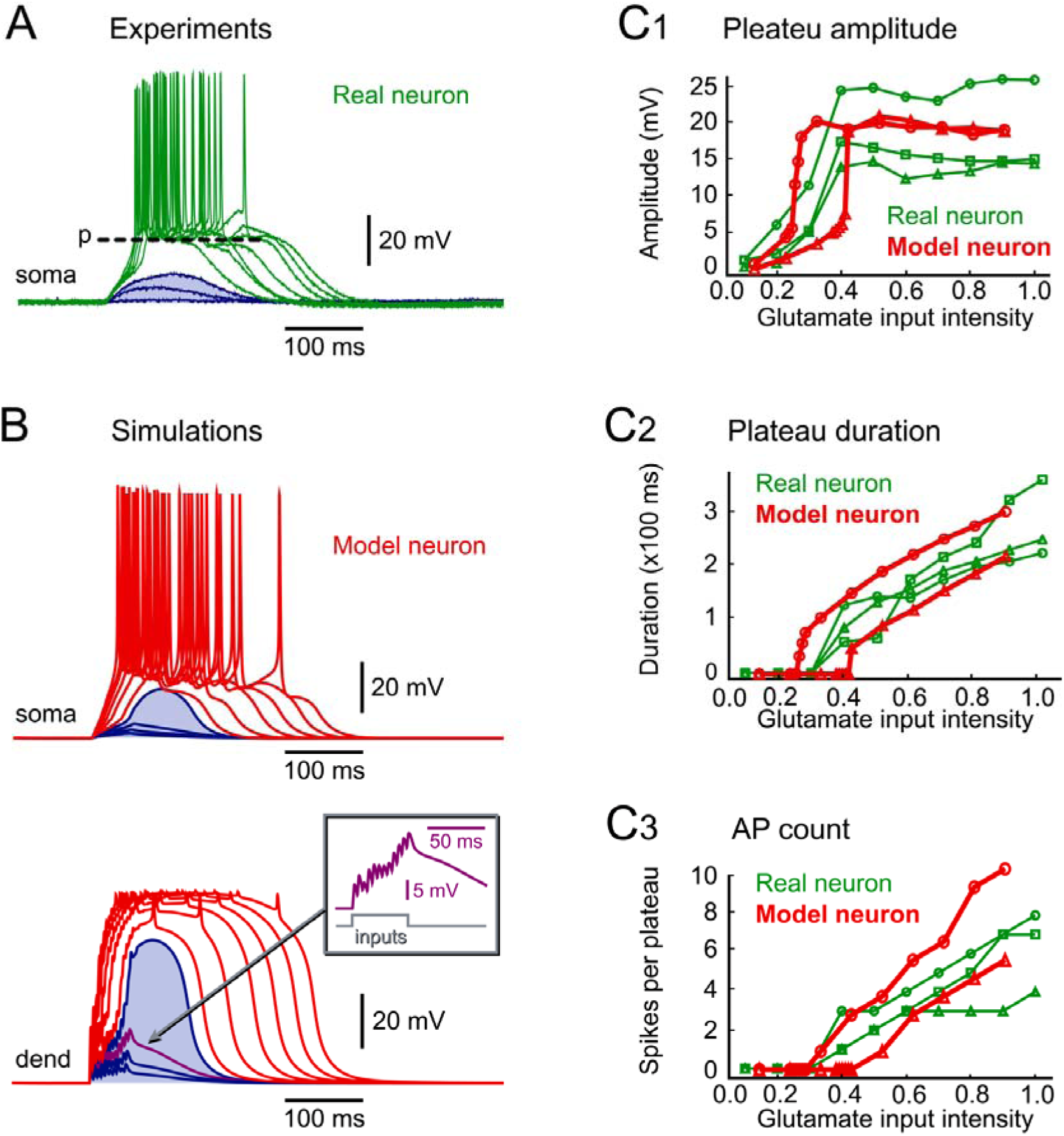
Somatic waveforms: experiments vs. simulations. **Varying levels of glutamatergic input in experiments and model. (A)** Somatic whole cell recordings in a layer 5 pyramidal neuron. Glutamate microiontophoresis was applied on one basal branch ∼90 μm away from the soma. The intensity of the glutamate iontophoretic current was increased gradually in equal steps. In this and all the remaining panels of the figure, subthreshold membrane responses are colored blue. **(B)** Computational model of simultaneous somatic (soma) and dendritic (dend) voltage recordings in response to glutamate application on one basal dendrite. Inset: Rectangle marks the time window for arrival of glutamatergic inputs on dendrite. Distance from the soma = 110 μm. The NMDAr mechanism here is based on Destexe et al., 1994 (our Model 1). The Model 2 data (employing the Major et al., 2008 NMDAr mechanism) is shown in **Suppl. Fig. S1B. (C**_**1-3**_) Numerical analysis of experimental data obtained in 3 real neurons (green) and 2 model neurons (red).

We built two models based on two existing mechanisms for NMDAR conductances. Our “Model 1” employed the Destexhe *et al*., 1994 NMDAR model, which mimicked our electrophysiological recordings with Type 1 morphology of the plateau collapse – gradual fall-off (Suppl. Fig. S1*A2*).

https://doi.org/10.6084/m9.figshare.11301020.v1

Our “Model 2” employed a more complex triple-exponential conductance model by Major *et al*., 2008, and was able to match Type 2 morphology of the plateau collapse – abrupt drop (Suppl. Fig. S1*B2*). Both Model 1 and Model 2 matched experimental plateau potential amplitudes, durations, and AP firing, all of which were similar for both Type 1 and Type 2 plateau morphologies (Suppl. Fig. S1, *C* and *D*). Performance of the two models was also similar, except for a difference in glutamate threshold. The Major *et al* NMDAR model (Model 2) required stronger synaptic activation for the NMDA current to become regenerative (e.g. spike).

Although voltage-sensitive dyes report membrane potential changes with microsecond precision, with an optical signal directly proportional to membrane voltage, they cannot give precise voltage values for the dendritic plateau (in millivolts), because dendritic optical signals cannot be calibrated using the somatic patch electrode; explained in ref. (Antic, 2003). Therefore, we used the simulation results to estimate plateau amplitudes at the site or their origin, in distal segments of thin dendritic branches. Modeling exercises predict that the plateau phase of the dendritic plateau potential is on average 57.6 ± 5.5 mV above resting membrane potential, or in the absolute range: −21 to −10 mV (24 stimulus locations 70–180 μm from soma center; 8 basal dendrites; both NMDAR models).

Somatic plateau durations were strongly correlated to dendritic plateau durations, both experimentally and in simulation (Fig. 4, *A* and *B*; R^2^=0.997 for simulation; R^2^=0.983 for experiment). While plateau durations (half-widths) were identical at all dendritic segments, the peak plateau amplitudes varied along the stimulated dendrite, being highest at the glutamate input site (Fig. 4*C*; input site at 85–115 μm). Plateau amplitude attenuated with distance from the input site with greater attenuation towards the soma and lesser attenuation towards the sealed end of the dendrite (Fig. 4*D*). bAP amplitudes (measured from plateau to peak, Fig. 4*C*, inset), on the other hand, were highest near the soma and attenuated to less than 5 mV at distal dendritic segments experiencing plateau potential (Fig. 4*E*, >100 μm). The attenuation of the bAP amplitudes was intensified around the glutamate input site (Fig. 4*E*, gray rectangle) suggesting a shunting effect imposed by the glutamate-activated dendritic conductances.

**Fig. 4.**
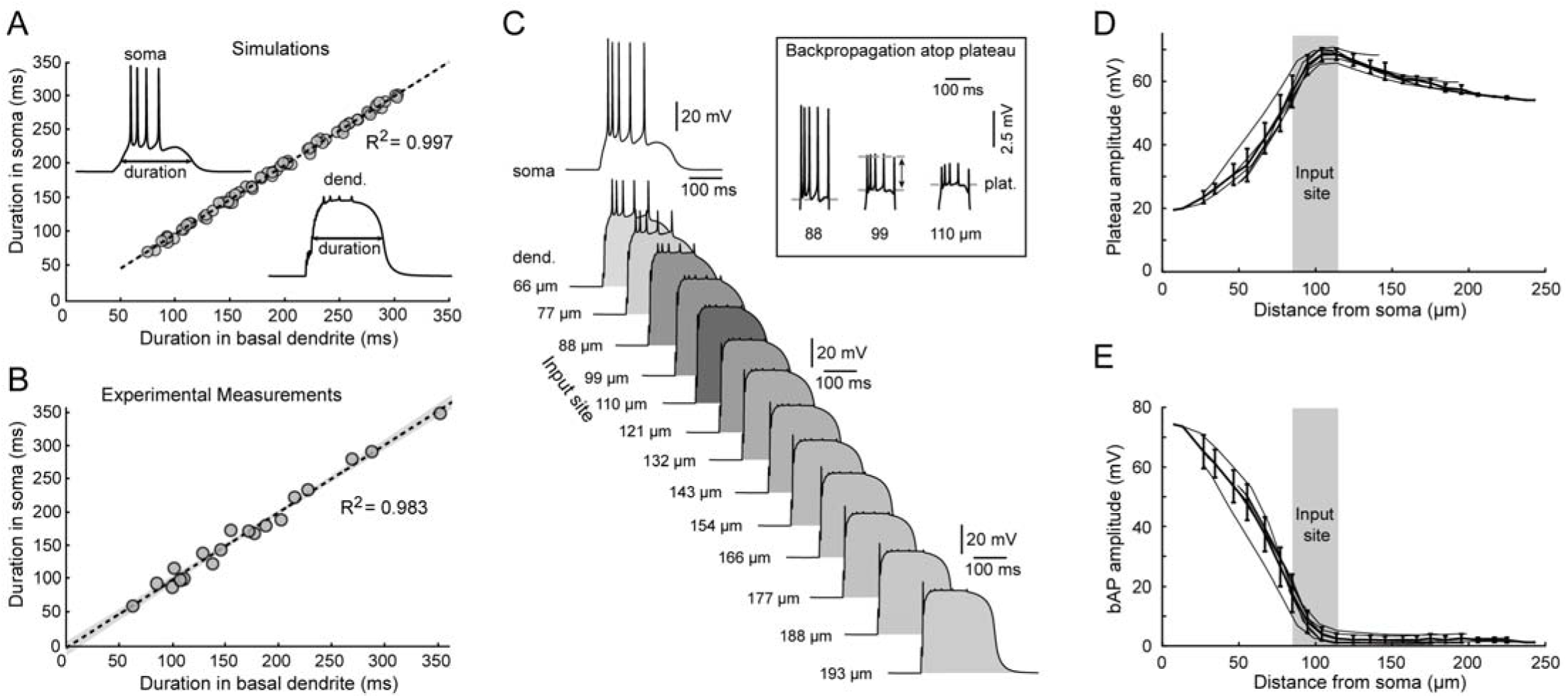
Plateau duration (half-width), plateau amplitude & “bAP on plateau”. **Voltage waveforms in soma and dendrite are strongly correlated**. **(A)** Comparisons between dendritic and somatic voltage transients obtained in the model simulations. n = 70 sweeps in 6 model basal branches. Insets depict the method used for measuring plateau amplitude and duration in dendrite and soma. **(B)** Experimental measurements in rat brain slices using simultaneous recordings of voltage waveforms in dendrite (voltage imaging) and soma (whole-cell). n = 19 sweeps from 5 pyramidal neurons. Linear fitting by ordinary least squares regression. The 95% confidence interval is marked by gray shading. **(C)** Computer simulation. Dendritic voltage waveforms obtained simultaneously from 13 locations along basal dendrite and cell body (soma). Inset: bAPs emerge from the plateau phase. Horizontal line “p” marks the amplitude of the plateau phase. Two-sided arrow indicates method used for measuring “AP amplitude above plateau”. **(D)** Dendritic plateau amplitude as function of distance from cell body. **(E)** Amplitude of bAPs above plateau level, as function of distance from cell body.

#### III The effect of input location on somatic depolarization amplitude

In published electrophysiological experiments (Fig. 5*A*), larger somatic plateau amplitude was associated with dendritic stimulation closer to the soma (Major et al., 2008). This experiment was reproduced in our model neuron (Fig. 5*B*), and showed distance fall-off similar to Major *et al* data (compare Fig. 5*A* vs Fig. 5*C*). Inverting the axes for the averages of the model data (Fig. 5*C*) allowed us to estimate the distance from the dendritic initiation site (in μm) as a function of somatic plateau amplitude (“amp” in mV):

**Fig. 5.**
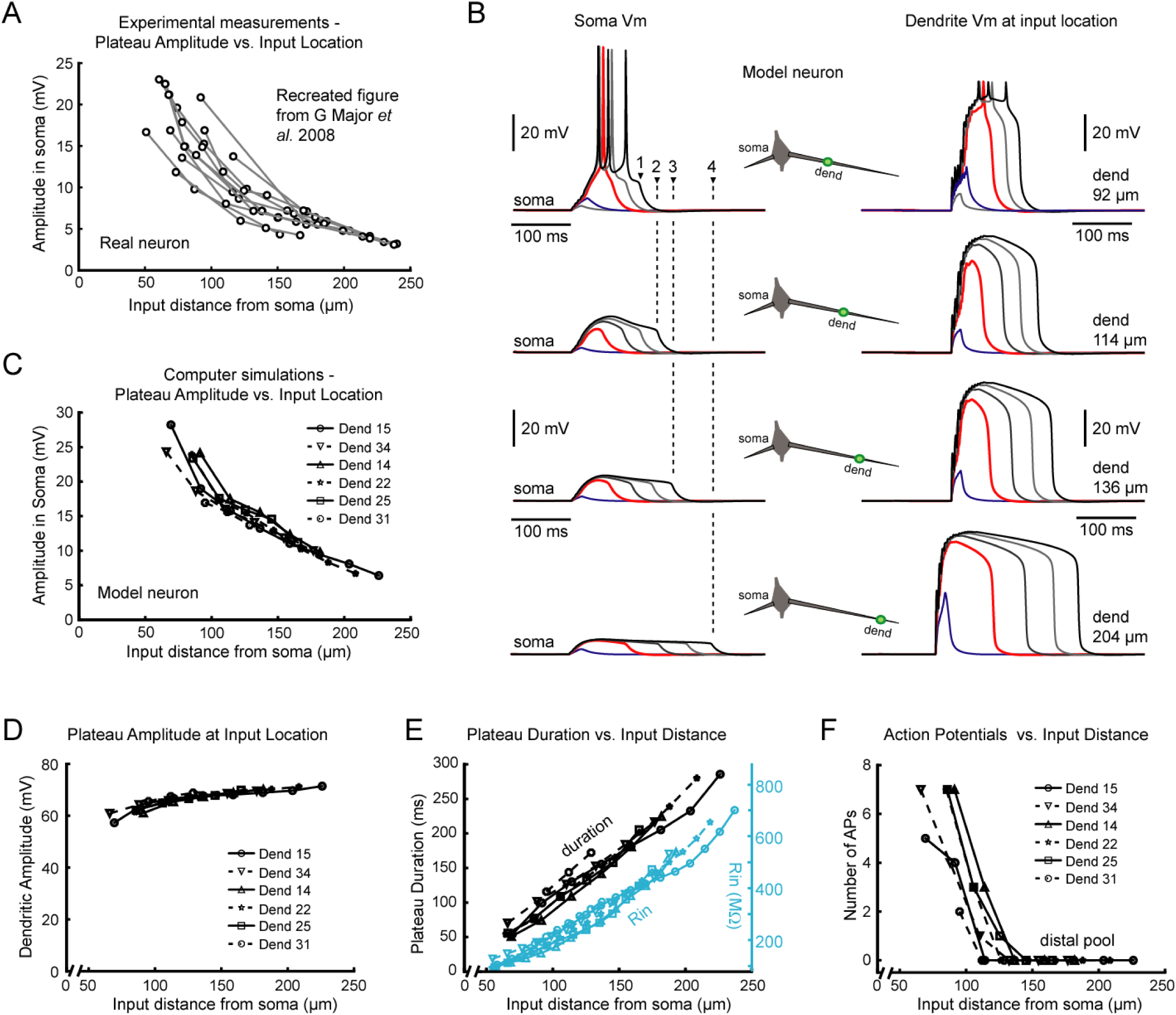
Glutamate input location influences: somatic plateau amplitude, plateau duration, and # of APs. **The impact of the input location**. **(A)** *Major et al (2008)* delivered glutamate microiontophoresis on the basal branches at various distances from the cell body and they measured membrane potential changes in the soma. The amplitude of the plateau potential in the soma is plotted against the distance from the cell body – recreated from *Major et al*., *2008*. **(B)** Computer simulation of the experiment described in *A*. Results obtained with our “Model 2” (Major et al., 2008) are displayed – simultaneous voltage waveforms in dendrite (at input location) and soma. The precise locations of the glutamate inputs on basal dendrite are annotated above each dendritic trace and expressed as a distance from the cell body in micrometers. Blue traces: sub-threshold depolarizations; Red: first suprathreshold, local regenerative potentials”. Vertical dashed lines mark 4 different plateau durations obtained by stimulating basal dendrite at 4 locations – fixed glutamate input intensity. The results of Model 1 (utilizing the Destexe et al. (1994) membrane mechanism for NMDAr channels) are shown in **Suppl. Fig. S2B**. **(C)** Same experimental outline as in *A*, except experiment was performed in simulations. Data quantifications of these simulation experiments are plotted. **(D)** In model neuron, the local amplitude of the dendritic plateau, measured at each glutamate stimulation site, is plotted versus the distance of that stimulation site from the cell body. **(E)** The local duration (half-width) of the dendritic plateau at the glutamate stimulation site is plotted as a function of distance from the cell body. Blue data points depict local dendritic R_in_. **(F)** Distal glutamatergic inputs (distal pool) generate dendritic plateau potentials, which fail to trigger somatic APs.

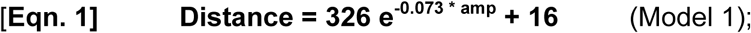

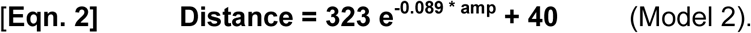

For example, the somatic plateau amplitude of ∼25 mV (Fig. 3*A*) would indicate that glutamatergic activation was received ∼68 μm away from the cell body, while amplitude 14 mV (Suppl. Fig. S1*B1*) would indicate a ∼128 μm distance. Overall estimated distance for the 15 cells studied ranged from 69 to 147 μm (average: 103 ± 19 μm).

Glutamate activation location only slightly affected dendritic spike amplitude at the stimulation location (Fig. 5*D*). This relatively subtle effect is explained by noting that plateau depolarizations are near the reversal potential of NMDAR voltage sensitivity, a ceiling effect. Location did have a substantial effect on plateau duration, an effect that could be seen in the soma as well as at the stimulus location, Fig. 5*B* (dashed vertical lines) and Fig. 5*E* (black markers). Duration effects can be explained by the gradual increase in input impedance (R_in_) from proximal to distal, seen due to the sealed end of the dendrite (R_in_ in Fig. 5*E*, blue markers).

Dendritic plateaus, propagating from dendrite to soma, often reach the threshold for AP initiation (Fig. 3*B*). The number of somatic APs depended on the glutamate input’s distance from the cell body (Fig. 5*F*). In each model dendrite, we found a similar distance, 128 ± 12 μm (n=6 dendrites; both models), beyond which distal inputs were subthreshold for somatic AP generation (Fig. 5*F*, distal pool). Presence or absence of APs atop the plateau phase is another way for estimating the distance of the strong glutamatergic input onto a basal dendrite. Inputs more distal than ∼130 μm typically produce spikeless plateau depolarizations in the cell body (Fig. 5*F*, distal pool). Although distal plateaus failed to drive AP initiation on their own, they caused sustained depolarizations of the cell body in the range of 10 to 20 mV. In the next series of experiments, we asked whether dendritic plateau potentials change the dynamics of the membrane response in the cell body.

#### IV Dendritic plateau potentials change global electrical properties of the neuron

To test the impact of dendritic plateau potentials on the overall neuronal membrane properties, input impedance (R_in_) and membrane time constant (TAU), rectangular current pulses (test pulses) were injected into the cell body of a model neuron, while glutamatergic input was delivered in the mid segment of one basal dendrite (Fig. 6*A1*, glut.). Simulations were performed in models with sodium channel blockade, mimicking treatment with TTX. Two test pulses of identical characteristics were delivered, before and during a glutamate-mediated plateau potential (Fig. 6*A1*). The cell body response to a rectangular current pulse underwent drastic changes in both amplitude (Fig. 6*A2*) and dynamics (Fig. 6*A3*). A decrease in steady state amplitude (Δ*Vm*) suggested that during a dendritic plateau potential, the cell body of the neuron was in a state of a lower R_in_. Furthermore, the neuronal membrane response was faster (Fig. 6*A3*, compare TAU-d vs. TAU-b). During a plateau, it took less time for the test-pulse-evoked voltage transient to reach 63% of its maximal amplitude (Fig. 6*A3*, 63%). That is, during dendritic plateau potential, the somatic TAU (TAU-d) is markedly shorter than the TAU measured before plateau onset (TAU-b) in the same neuron.

**Fig. 6.**
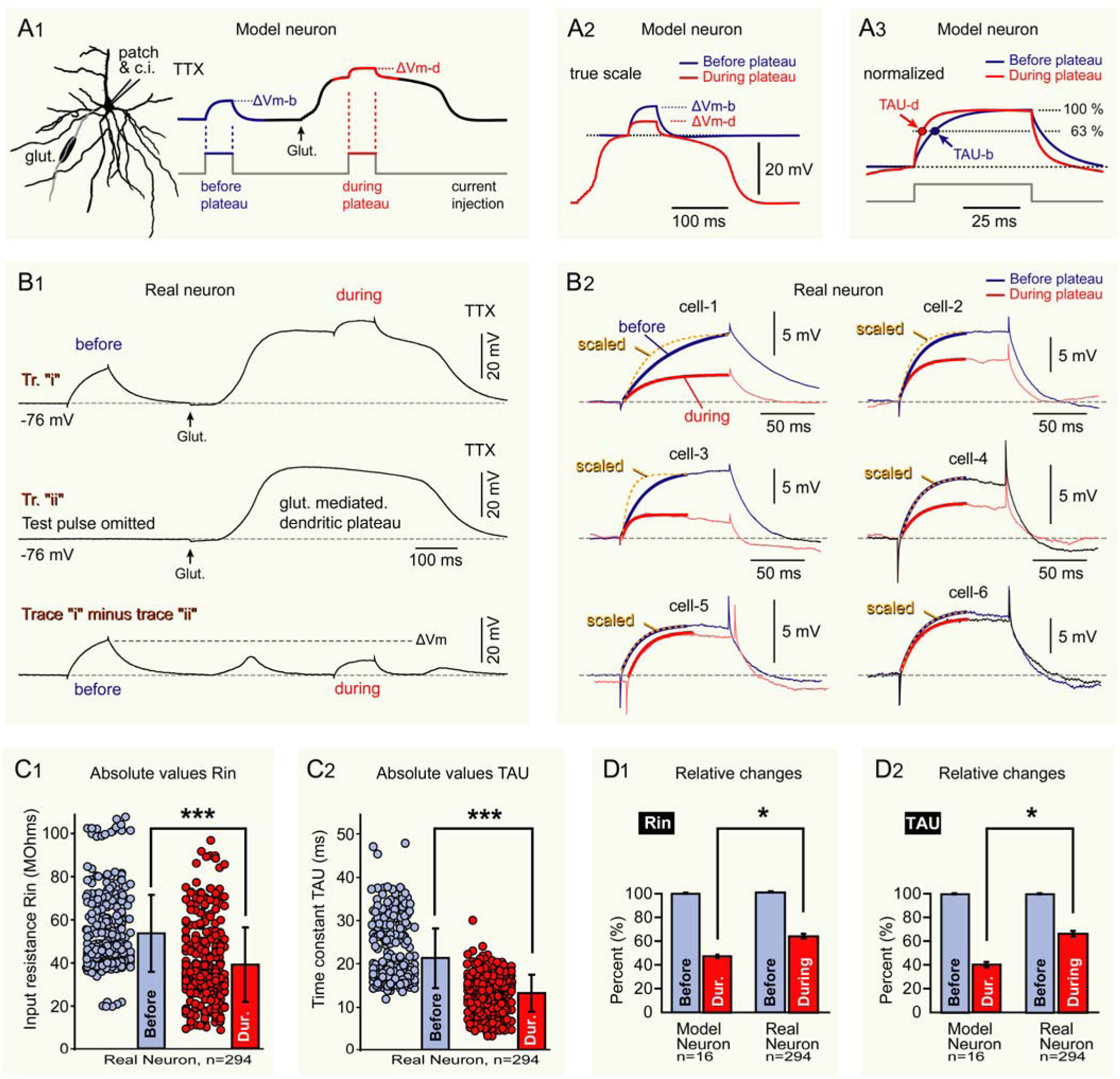
Plateau induced changes on the membrane charging curve. **The cell body TAU and R_in_ are affected by dendritic plateau potentials**. **(A**_**1**_) Basilar dendritic tree of the model cell. Dendritic segment (66–132 μm) is receiving glutamate inputs. Computer simulation of dendritic plateau potential measured at soma. An identical depolarizing current pulse was injected into the soma before and during the plateau potential (current injection). **(A**_**2**_) A1 responses superimposed before (blue) and during plateau (red); amplitude during plateau (dVm-d) is smaller than before plateau (dVm-b). (**A**_**3**_) A1 responses superimposed and normalized to TAU-b. **(B**_**1**_) Whole-cell recordings in TTX with 5 ms glutamate pulse at dendritic location 90 μm from soma. Test pulses for testing R_in_ and TAU were attained by somatic current injection, “before” and “during” plateau, as in the model (5 sweeps average for each trace). In trial “ii” test pulses were omitted to reveal the waveform of the underlying plateau. Subtraction “i” minus “ii” at bottom shows responses to test pulses, free from plateau induced wobbles in the baseline. **(B**_**2**_) Comparisons of evoked responses before and during plateau. 6 cells are displayed to show cell-to-cell variability. Scaling was used to allow comparison of time constants. **(C)** Raw values. R_in_ and TAU both decrease during plateau potential (mean ± standard deviation; n=294 trials in 18 dendrites of 8 neurons p<0.0001 ***). **(D)** Relative values (During/Before). Comparison of model (n=16 trials obtained by stimulation of 16 dendrites, in 2 model neurons) with experiment from **C** using relative values (* indicates p<0.01).

These model predictions (Fig. 6*A1*) were then tested experimentally (Fig. 6*B1*) using an identical paradigm, in the presence of sodium channel blocker TTX, 1 μM. We compared two types of traces recorded from the same cell: [i] traces with glutamatergic stimulation paired with test pulses (Fig. 6*B1*, “i”); and [ii] traces with test pulses omitted, i.e. glutamatergic stimulation only (Fig. 6*B1*, “ii”). The contour of the glutamate-induced plateau, unaltered by test pulse (test pulse omitted), was used to eliminate undulations in the baseline caused by the underlying plateau (Fig. 6*B1, Trace “i” minus trace “ii”*).

During the dendritic plateau potential, test-pulse-induced somatic voltage transients were smaller in amplitude (ΔVm) and faster to rise (shorter TAU), compared to the same test pulse performed on the same neuron just before the plateau onset (Fig. 6*B2*, compare traces *during* vs. *before*). In the majority of cells, *dVm* and *TAU* decreased significantly during the plateau (Fig. 6*B2, cell-1* to *cell-4*). However, in some trials these changes were more subtle (Fig. 6*B2, cell-5* & *cell-6*). Average R_in_ (n=294 traces in 18 dendrites belonging to 8 neurons) was 53.6±1.0 MΩ before plateau, and 39.1±0.9 MΩ during plateau (Fig. 6*C1*, p<0.0001, unpaired student’s t-test). Average TAU was 21.3±0.4 ms before plateau, 13.2±0.2 ms during plateau (Fig. 6*C2*, p<0.0001, unpaired student’s t-test).

Plateau-induced changes of R_in_ were predicted by the model (Fig. 6*A*): decrease of R_in_ and TAU values during plateau. We analyzed the magnitude of the plateau-induced change in model and real neuron. In each trace of the current data set we measured membrane response of the same neuron (model and real) before and during glutamate-evoked plateau potential. This allowed us to calculate the ratio *During/Before* for each trace in both Model and Real neurons of this study. The ratio “*During/Before*” is a measure of a relative decrease in R_in_ or TAU due to underlying plateau potential. R_in_ During/Before simulation: 48±1.5 % (mean ± sem; n = 16 trial locations, 16 dendrites, 2 model cells, Fig. 6*D1*, Model Neuron). R_in_ During/Before experiment: 64.5±1.4 % of the (n = 294 trials, 18 dendrites, 8 cells, Fig. 6D1, Real Neuron; *, p<0.01). TAU During/Before simulation: 39.6 ± 0.6 % (Fig. 6*D2*, Model Neuron), or 65.9 ± 1.7 % in real neurons (Fig. 6*D2*, Real Neuron; *, p<0.01). In ∼13% of experimental dendritic locations (n = 39 out of 294 recordings), we found a different result with respect to TAU, with TAU-d equal or greater than TAU-b (Fig. 6*B2, cell-5* & *cell-6*). This anomaly was in one part due to some distal plateaus having a small amplitude in the cell body (see below), and partially an artifact of the difficulty of controlling baseline voltage during the plateau (Fig. 6*B1*, note a slow decline of voltage during plateau phase), which would greatly alter estimation of TAU by exponential fitting. Including the outliers, the degree of shortening was less than predicted (Fig. 6*D2*, *).

Using sinusoidal current injections into the cell body we found that neuronal impedance decreases during plateau potential (Suppl. Fig. S3). The effect was strongest at low frequencies (5 and 10 Hz), and negligible at stimulus frequencies greater than 75 Hz (Suppl. Figs. S4 - S7).

We have shown that glutamatergic inputs arriving in proximal segments of basal dendrites, closer to the cell body, produce greater somatic depolarizations than input received in distal dendritic segments, far away from the cell body (Fig. 5). Next, we asked whether proximal dendritic inputs exert stronger influence on the dynamics of the somatic membrane response (Fig. 7*A1*). Using fixed input intensity, the membrane charging curve was faster when plateau potential was induced by proximal inputs (Fig. 7*A2*, inset, compare “*distal*” and “*proximal*” trace). By gradually changing the location of glutamatergic input along model basal dendrites (n = 6 dendrites) and measuring soma TAU on each trial, we found that plateau-induced shortening of the soma TAU in model neurons strongly depended on the distance between the cell body and glutamate input site - proximal inputs exerted more prominent shortening of the soma TAU than distal inputs (Fig. 7, *B1* and *B2*).

**Fig. 7.**
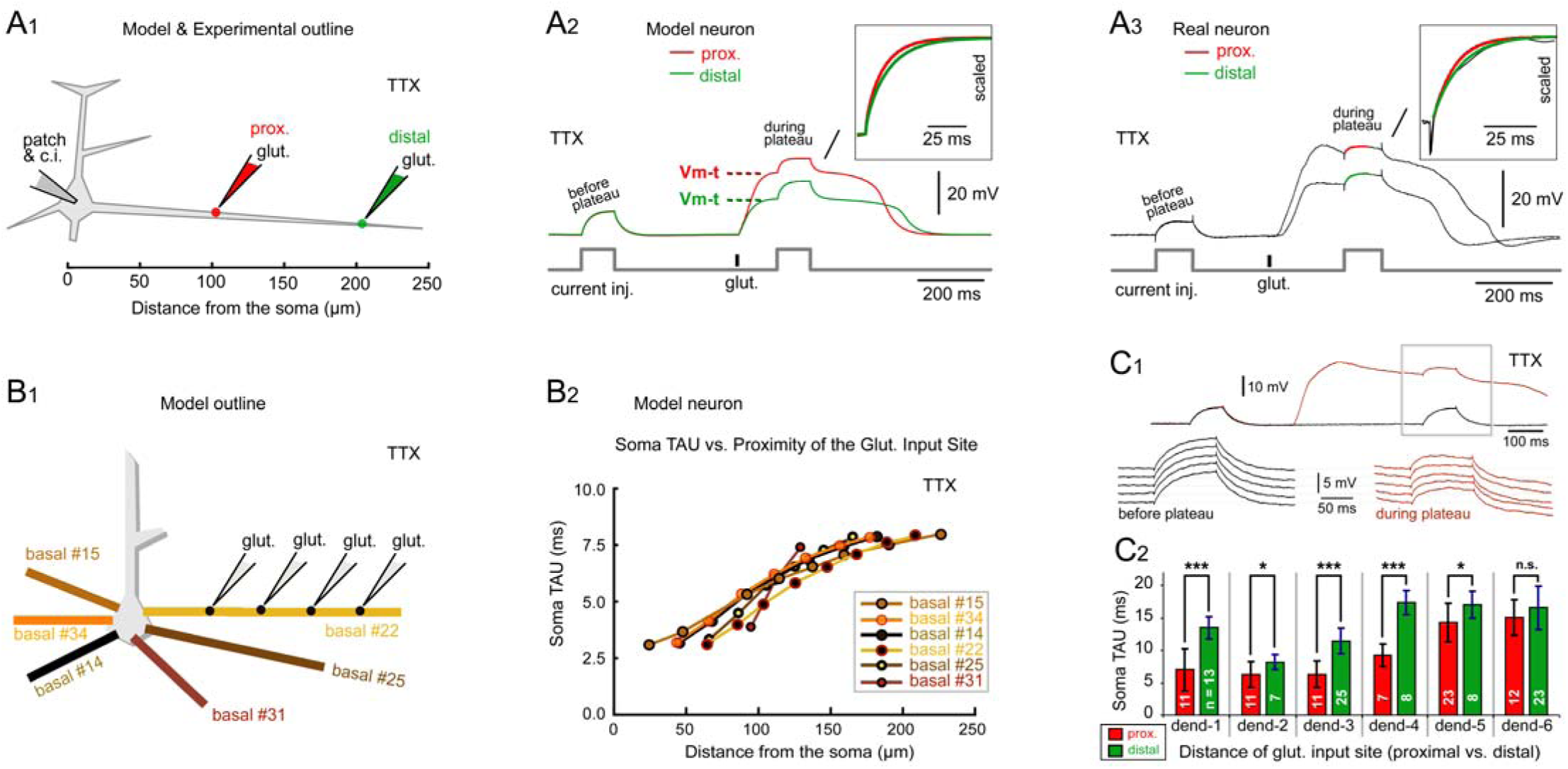
The cell body charging curve as function of “*proximal”* vs. “*distal”* input locations. **Position of glutamatergic input on basal dendrite determines the magnitude of the plateau-induced changes in the somatic TAU**. **(A**_**1**_) Experimental outline in model and experiment: in the presence of TTX, glutamate input of fixed intensity delivered at two different locations along a basal dendrite. **(A**_**2**_) In model, plateau amplitude at cell body has greater amplitude and faster rise when triggered from proximal location (red) compared to distal location (green; inset: comparison of charging curves). Vm-t marks the membrane voltage at which the test pulse “during plateau” begins. **(A**_**3**_) Experiment verifies model prediction. **(B**_**1**_) Schematic of input location shifting on a basal branch with input intensity fixed. Multiple dendrites are examined. **(B**_**2**_) TAU increased with increased distance of the glutamate input site, for all 6 dendrites tested. **(C**_**1**_) In real neurons, multiple traces from one glutamate stimulation site were recorded with glutamatergic stimulation ON (brown) or OFF (black). A blow up of membrane responses before and during plateau; multiple repetitions. **(C**_**2**_) Comparison of TAU between distally and proximally delivered, fixed-intensity-glutamatergic input on the same dendrite (distal site 40 – 110 μm away from proximal). Error bars are standard deviations; (***) p<0.01; (*) p<0.05; (n.s.) “not significant”).

Experiments supported prediction: plateau-induced changes of cell body R_in_ and TAU are more pronounced when dendritic plateaus were more proximally. The model prediction was tested using TTX to prevent any AP firing (Fig. 7*A1*, schematic). At each input location we recorded multiple repetitions, with and without glutamate stimulus (Fig. 7*C1*). With distances between proximal and distal input site of 40 – 110 μm, membrane charging was faster when the plateau potential was induced by proximal dendritic inputs. More specifically, in 5 out of 6 dendrites stimulated at two locations, the soma TAU were significantly shorter in proximal versus distal paradigm (Fig. 7*C2*), as predicted by model (Fig. 7*B2*).

#### V Voltage-induced changes in R_in_ and TAU

Although our working hypothesis states that plateau-induced changes in TAU are due to massive dendritic conductances, greater somatic depolarization with more proximal input (Fig. 5) might also explain the greater effects of proximal inputs on R_in_ and TAU. We therefore tested R_in_ and TAU while controlling somatic voltage by either 1s current clamp (Fig. 8*A*, n=22 neurons), or dendritic plateau potential (Fig. 8*B*, n = 8 neurons), in the presence of TTX. A test pulse was delivered in the middle of the “voltage-setting pulse” (Fig. 8*A1*, test pulse). The neuronal membrane potential, Vm-t, obtained just before the arrival of the test pulse, served as an independent variable in this experiment, whereas R_in_ and TAU are dependent variables. Measures were normalized against values obtained in the same neuron at RMP, showing shallow hyperbolic distributions across the range of membrane voltages, “Vm-t”, (Fig. 8, *A2* and *A3*). Too much depolarization, or too much hyperpolarization, decreased both R_in_ and TAU.

**Figure 8.**
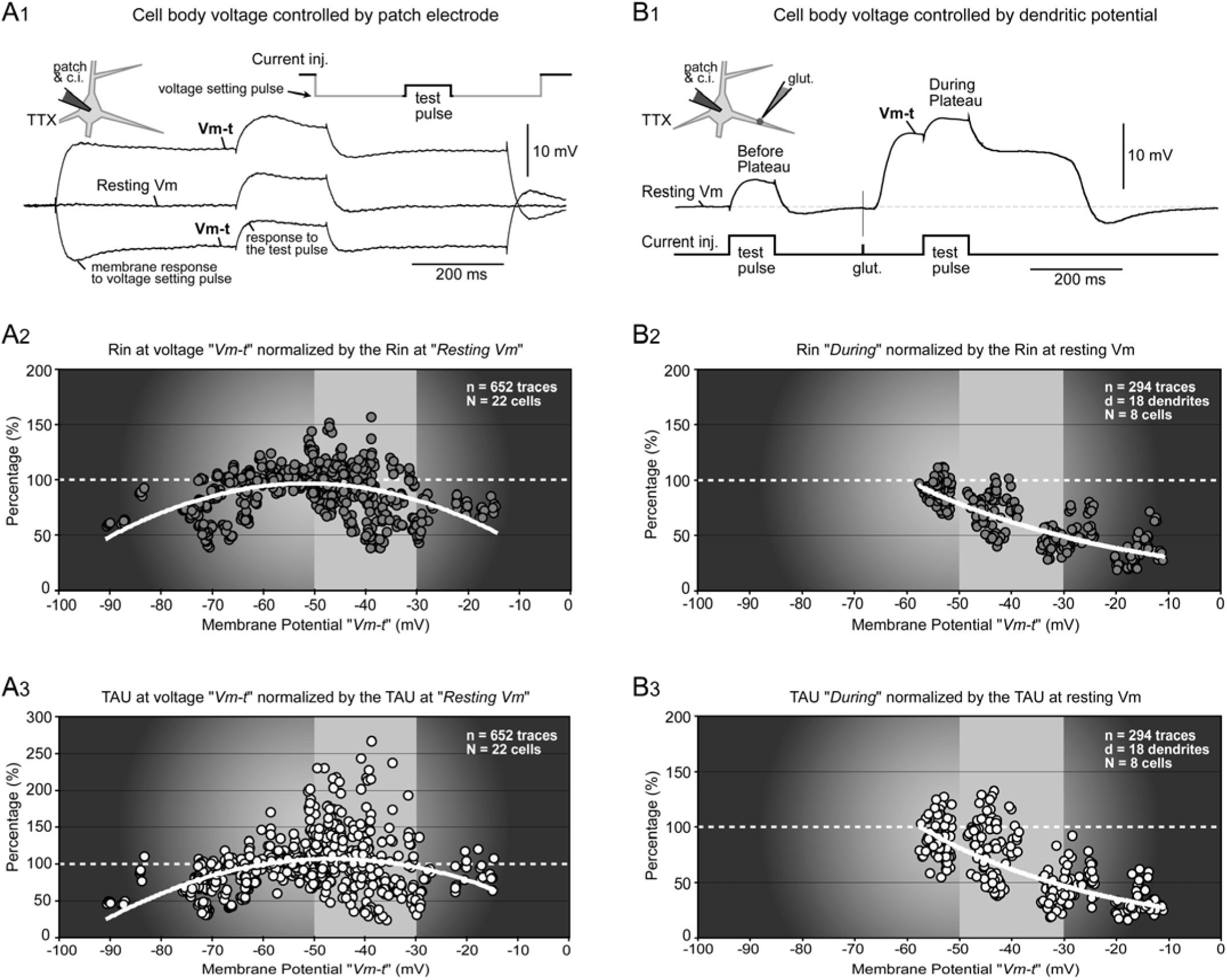
The effect of membrane potential on neuronal Rin or TAU. **Dendritic plateau potential alteration of somatic TAU and Rin is not fully explained by effect on Vm**. Experiments in TTX (A1) Vm with current injection (“voltage setting pulse” Inset). Additional current pulse (Inset: “test pulse”) was used to measure R_in_ and TAU. “Vm-t” is value at start of test pulse. (A2) Soma R_in_ normalized by R_in_ at resting membrane potential (RMP), plotted against the Vm-t. (A3) Soma TAU normalized by TAU at RMP, plotted against the Vm-t. (B1) Glutamate evoked plateau potential with test pulse. (B2) Soma R_in_ normalized by R_in_ at RMP, plotted against the Vm-t. (B3) Soma TAU normalized by TAU at RMP, plotted against the Vm-t. Second order polynomial fits without intercept. The 2 conditions (A and B) differ markedly from −50 to −30 mV (light gray).

Effects were more marked when using voltage change via dendritic plateau potentials (Fig. 8*B1*, n=8 neurons). R_in_ and TAU both still decreased with membrane potential increases (Fig. 8, *B2* and *B3*). However, plateaus produced more consistent reductions in both measures, particularly in the range of −50 to −30 mV (Fig 8, highlighted area). The more pronounced effects with plateaus indicate that voltage alone does not explain the R_in_ and TAU changes. Instead, glutamate-mediated dendritic plateau potentials affect somatic R_in_ and TAU through combined effects of both voltage and conductance changes.

#### VI Plateau potentials and synaptic integration

Plateau depolarization brings the somatic membrane closer to firing threshold (Fig. 5C), which may enhance the efficacy of EPSPs towards initiation of APs. Faster charging of the somatic membrane (lower TAU), may further enhance the efficacy of EPSPs in producing a spike. However, reduction in R_in_, and reduction in driving force, will reduce EPSP amplitude and reduce this boost. To assess these countervailing influences, we simulated integration of identical sets of spatially-distributed EPSP barrages arriving on multiple dendrites (Fig. 9) in the presence, or absence of dendritic plateau potential occurring in a single basal dendrite (Fig. 9A, dendrite marked by “*Plateau*”). An EPSP barrage that was subthreshold before the plateau (Fig. 9*B*, “*EPSPs before plateau”*), caused AP firing when it arrived during the plateau (Fig. 9*B*, “*EPSP-evoked AP*”), demonstrating that “a spikeless” plateau will enhance the efficacy of synaptic integration.

**Fig. 9.**
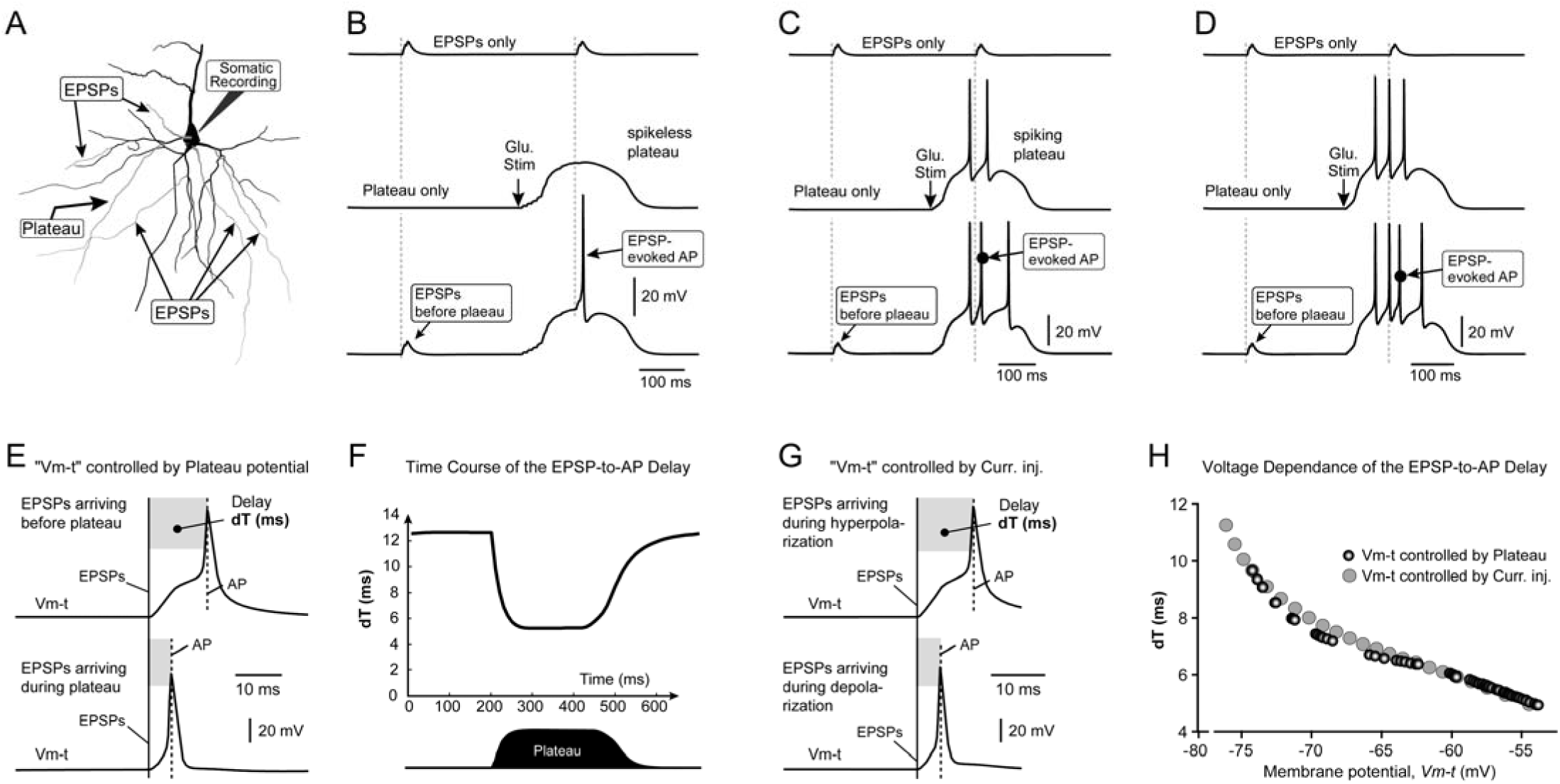
The impact of dendritic plateau potential on axonal AP Initiation. **A dendritic plateau potential occurring in one basal dendrite influences the somatic integration of EPSPs arriving on other dendrites. (A)** Basilar dendritic tree of the model cell. Plateau was produced in 1 branch (“plateau”). Individual EPSPs were received at 5 locations marked by EPSP-arrows. **(B)** Spikeless plateau is paired with EPSPs. Top trace (*EPSP only*): An identical barrage of EPSPs was delivered twice, causing 2 EPSP events. Middle trace (*Plateau only*): Glutamatergic stimulation of one basal dendrite produced a somatic plateau subthreshold for axonal AP initiation (spikeless plateau). Bottom trace: pairing of the stimulation paradigms used in the top and middle trace. **(C)** “Spiking plateau” (plateau accompanied by somatic APs) is paired with EPSPs. Intercalated spike is marked by black dot and label “*EPSP-evoked AP*”. **(D)** The same modeling experiment as in *C*, except an even stronger plateau, with more accompanying APs, was used for pairing. **(E)** Upper trace: Strong EPSP barrage, capable of triggering AP in the absence of plateau. Lower trace: Pairing of the EPSP barrage with a spikeless plateau. Full vertical line marks the onset of the EPSP barrage (EPSPs). Dashed vertical line marks the peak of the EPSP-evoked AP (AP). Gray box marks a time delay between the onset of EPSP barrage and the AP peak (dT). “Vm-t”, voltage just prior to arrival of EPSPs. **(F)** dT is reduced during plateau. **(G)** Voltage controlled by current injection --compare with *E*. A more depolarized Vm-t produces AP sooner. **(H)** The voltage dependence of the dT is similar with and without dendritic plateau potential.

We also investigated the effects of plateaus crowned with AP (“spiking plateaus”) on EPSP barrages (Fig. 9C), demonstrating a distinct EPSP–induced AP can also be generated during spiking plateaus (Fig. 9C, “*EPSP-evoked AP*”). The timing of the additional spike (“*EPSP-evoked AP*”) was closely tied to the EPSP onset (dashed vertical line), regardless of the number of APs riding on the plateau phase (Fig. 9, *C* and *D*). Hence, both “spikeless plateaus” (Fig. 9B) and “spiking plateaus” (Fig. 9, *C* and *D*) increased the capacity of cortical neurons to respond to afferent EPSPs by generation of new APs (“*EPSP-evoked AP*”).

The amount of time (temporal delay) from the onset of EPSP barrage to the AP peak (dT) can be used as a measure of the neuronal responsiveness to incoming afferent inputs. Shorter dT would indicate neuronal states of greater excitability. In these series of simulations, the dendritic plateau potential was kept at a fixed time delay from the beginning of the trace (e.g. time point 200 ms). In repeated simulation trials, we systematically changed the timing of the EPSP barrage (from 0 to 600 ms), while keeping the plateau fixed at 200 ms. The EPSP-to-AP temporal delay (dT) was reduced from ∼12 ms before plateau to ∼6 ms during plateau (Fig. 9E). Time interval dT precisely followed the contours of the plateau voltage waveform (Fig. 9F), suggesting a strong impact of dendritic plateau potentials on the process of synaptic integration in the cell body. During the plateau potential, the interval between EPSP onset and AP peak shortened, with degree of shortening proportional to depolarization, demonstrating greater responsiveness to incoming EPSPs.

The plateau induced changes in dT (Fig. 9F) could be due to either depolarization of membrane, or shortening of TAU (Fig. 6), or both. To distinguish the impact of membrane voltage alone, we designed an approach in which we compare the behavior of dT, while the cell body voltage was controlled by either patch pipette (Fig. 9G, Current injection) or dendritic plateau potential (Fig. 9E). More depolarization produced shorter time intervals to AP in response to a standard EPSP barrage (Fig. 9, *G* and *H*). Comparing the effects of somatic current injection (Fig. 9H, light gray circles) to dendritic plateau potential (Fig. 9H, dark gray circles) at the same membrane voltage, demonstrated only slight additional effect of time constant change in decreasing dT.

## Discussion

We used optical imaging to characterize voltage waveforms of glutamate-mediated plateau potentials occurring in basal and oblique dendrites of neocortical layer 5 pyramidal neurons. By employing a gradually increasing intensity of the focal glutamatergic input, we documented the transitions between subthreshold to threshold regenerative local potentials, occurring at the dendritic site of initiation. Previously, subthreshold-to-threshold transitions of membrane potential occurring in a thin basal dendrite were inferred indirectly from the somatic patch electrode recordings (Schiller et al., 2000; Losonczy et al., 2008; Major et al., 2008; Takahashi and Magee, 2009; Augustinaite et al., 2014). The sets of voltage waveforms, obtained at different levels of excitatory input, and simultaneously recorded at dendrite and soma (present study), provided constraints for simulation of a cortical pyramidal neuron. The modeling data obtained with the detailed model (realistic morphology + uneven spatial distributions of membrane conductances) demonstrated that dendritic plateau potentials changed the state of pyramidal cells in a profound way, with implications for neuronal network information processing.

During these dendritic plateau potentials, cell bodies of pyramidal neurons are placed in a depolarized state closer to the AP firing threshold. With this sustained depolarization state, the somatic membrane shows a notably faster capacitative charging in response to depolarizing currents (i.e. excitation). As a result of dendritic plateaus, pyramidal neurons are more responsive to afferent synaptic activity arriving anywhere on the complex dendritic tree. Barrages of EPSPs, which are subthreshold before the plateau, become more successful drivers of axo-somatic APs during the plateau. Even in spiking neurons, dendritic plateaus accelerate initiation of additional EPSP-induced APs. The time delay between the onset of EPSPs and the onset of EPSP-induced AP (dT) is notably shorter during the plateau. Faster EPSP-to-AP transition rates are expected to improve neuronal ability for “tuning” into fast rhythmic afferent synaptic and network activities. Dendritic plateau potentials move cortical pyramidal neurons from resting state (‘Down state like’) into a more excitable state (‘Up state like’); a sustained depolarized state during which afferent inputs are more effective and the transformation of afferent excitatory inputs into postsynaptic AP firing is faster.

### Dendritic voltage waveforms

Over the years, researchers have used somatic recordings to study initiation of local spikes in basal and oblique dendrites of pyramidal neurons. Within somatic recordings, they searched for “kinks” which may represent filtered and decremented versions of the original dendritic spike waveforms (Losonczy et al., 2008; Remy et al., 2009). A lot has been learned about the behavior and ionic composition of local dendritic spikes without ever showing what dendritic spikes look like in the thin dendrite (Losonczy et al., 2008; Remy et al., 2009). In the current study, we overcame the limitations of the earlier explorations by employing an optical imaging method capable of tracking membrane potential changes at submillisecond resolution (Short et al., 2017). We show that glutamate-mediated local dendritic spikes are complex waveforms comprised of several phases including a rapid rise, initial spikelet, plateau segment and abrupt collapse back to resting membrane potential (Fig. 1*C*).

In one prior study, researchers were able to patch basal dendrites with micropipettes of very high electrical resistance, resulting in dendritic whole-cell recordings with high series resistance (Nevian et al., 2007). They found that synaptic stimulations generate rectangular local dendritic potentials, while direct current injections generated local sodium spikes in some basal dendrites. Our current study explored several aspects of dendritic potentials which were not measurable in the prior study [1-5]. [1] We explored how gradual increase in glutamatergic input triggers local regenerative potentials in basal dendrite. The previous study did not explore basal dendrite voltage in response to gradually increasing input (but see ref. (Larkum et al., 2009) for graded glutamatergic input on apical tuft branches). [2] We showed that sodium spikelets were initiated by glutamatergic inputs. The previous study used direct current injection to activate sodium channels. [3] We showed that sodium spikelets precede the plateau phase and are responsible for the same kinks in the somatic recordings (Fig. 1*C*), previously correctly interpreted as dendritic spikes in thin branches (Losonczy et al., 2008; Remy et al., 2009). [4] We made recordings from two dendritic sites simultaneously (Fig. 1*D*) allowing for a better understanding of dendritic voltage maps. The previous studies ware restricted to one recording site per basal dendrite. [5] We described propagation of glutamate-evoked sodium spikelets and plateau potentials from dendrite to soma and simultaneous propagation of APs traveling from soma to dendrite. These three types of potentials meet in the mid segments of dendritic branches resulting in complex waveforms, which have not been identified previously by patch electrode recordings in basal dendrites (Nevian et al., 2007).

### Experimental constraints on the current model

Typically, computational models of CNS neurons are based on the voltage recordings from the cell body. Our model of a cortical pyramidal neuron was constrained by 4 sets of experimental data: (i) voltage waveforms obtained at the site of the glutamatergic input in distal basal dendrite, including initial sodium spikelet, fast rise, plateau phase and abrupt collapse of the plateau; (ii) a family of voltage traces describing dendritic membrane responses to gradually increasing intensity of glutamatergic stimulation (Fig. 1, *A* and *B*, ‘dend’); (iii) voltage waveforms of backpropagating action potentials in basal dendrites (Antic, 2003); and (iv) the change of bAP amplitude in response to drugs that block Na^+^ or K^+^ channels (Acker and Antic, 2009).

### Extrasynaptic NMDAR channels

Glutamatergic synapses are functionally clustered on dendrites; the neighboring synapses in one dendrite activate together more often than the synapses scattered on many branches (Larkum and Nevian, 2008; Kleindienst et al., 2011; Wilson et al., 2016). Repetitive clustered synaptic stimulation has been shown to overcome the ability of astrocytic processes to clear glutamate (Suzuki et al., 2008; Chalifoux and Carter, 2011), and can actually induce astrocytic glutamate release through reversal of astrocyte glutamate transporters (Carmignoto and Fellin, 2006). This would allow excitatory neurotransmitter to spill-over from synaptic clefts to activate extrasynaptic NMDA receptors between dendritic spines (Rusakov and Kullmann, 1998; Tovar and Westbrook, 1999; Harris and Pettit, 2007; De-Miguel and Fuxe, 2012; Petralia, 2012). During synaptically-evoked NMDA spikes (2 synaptic stimuli at 50 Hz), glutamate diffusion from synaptic clefts to extrasynaptic NMDA receptors on dendritic shafts has been detected using two-photon calcium imaging (Chalifoux and Carter, 2011).

Direct immunostaining has found a similar density of NMDARs at synaptic and extrasynaptic locations (Petralia et al., 2010). A kinetic study showed no major differences in dynamics between NMDARs at these two locations (Papouin and Oliet, 2014). This led us to set magnitudes of synaptic and extrasynaptic NMDA conductances at a 1:1 ratio, and to use the same membrane mechanism for both (Methods). We implemented a 5 ms time delay to account for time required for glutamate diffusion from synaptic cleft to dendritic shaft.

Our model demonstrated dendritic plateaus even without glutamate spillover. However, adding the activation of extrasynaptic NMDARs improved three cardinal features of dendritic plateau potentials (Fig. 2*C2*). Namely, activation of extrasynaptic NMDARs produces: [a] faster rising, [b] longer lasting, and [c] larger amplitude somatic plateau depolarizations. We speculate that the major difference between typical dendritic NMDA spikes, which last ∼50 ms (Schiller et al., 2000), and typical dendritic plateau potentials, which last 200 – 500 ms (Milojkovic et al., 2004; Suzuki et al., 2008), may lie in the extent to which a released glutamate is maintained around dendrite, due to a standstill in the astrocytic uptake, causing the activation of extrasynaptic NMDARs (Oikonomou et al., 2012).

### Plateaus last longer when triggered from distal dendritic segments

(Fig. 5*E*, “*duration*”). This effect is due to higher distal local effective R_in_ (Fig. 5*E*, “*R*_*in*_”). Consider two identical glutamate receptor current waveforms, one generated at proximal, and the other occurring at distal dendritic location. The waveform of the NMDA current interacts with the local R_in_ to generate a local voltage transient (ΔVm = R_in_ * I_NMDA_). In distal dendritic segments equipped with higher R_in_, the tail of the current waveform produces greater voltage deflections (for longer time) thus giving local plateau potentials longer durations. Our model predicts that distally positioned synaptic cluster would produce longer sustained depolarizations compared to the proximally positioned cluster. Therefore, strategic positioning of a distinct group of afferents on the most distal dendritic segments may have an impact on cortical processing of information (Antic et al., 2018).

### Spikeless plateaus begin in the middle of the basal dendrite

As glutamate input location is moved away from the soma, somatic plateau depolarization amplitude decreases rapidly (Milojkovic et al., 2004; Major et al., 2008; Augustinaite et al., 2014). This “distance-dependence” segregates roughly the basal dendrite into proximal and distal region -- only proximal dendritic segments will generate APs on the somatic plateau (Fig. 5*F*). Distal dendritic segments, on the other hand, produce spikeless depolarizations of the soma (Fig. 5*B*, 114 μm). We predict that this distal “no-AP-generation region” begins at 110-130 μm in most basal dendrites (Fig. 5*F*), consistent with ref. (Milojkovic et al., 2004) - their figure 1. The functional implications of having two context-defined groups of excitatory inputs segregated in two geographical sections of the same dendrite, “proximal drivers” (arriving in proximal dendrite) versus “distal drivers” (impinging on distal dendritic segments) are discussed in ref. (Jadi et al., 2014); while “retinothalamic” (proximal dendrite) versus “corticothalamic” (distal dendrite) input groups, and their biophysical interactions, are discussed in ref. (Augustinaite et al., 2014).

### Invasion of backpropagating APs into the plateau generating dendrite

An AP backpropagating from soma to dendrite maintains significant amplitude out to ∼150 μm (Fig. 2*B2*, black line). During a plateau, APs backpropagating from soma to dendrite are readily seen above the plateau at proximal, but not at distal locations (Fig. 4*C*), with bAP on plateau amplitude-decline abruptly occurring at 70-100 μm (Fig. 4*E*). Backpropagation of APs on plateau (from soma to dendrite) is limited by four factors: (i) lack of regenerative Na^+^ channel activation distally due to low channel density in dendrite; (ii) plateau depolarization-induced Na^+^ channel inactivation; (iii) plateau-depolarization-induced repolarizing current from various K^+^ channels in basal dendrites (Cai et al., 2004; Nevian et al., 2007; Acker and Antic, 2009); and (iv) shunting of the AP current through the large NMDAR conductance at the glutamate input site (Fig. 4*E, input site*).

### Distortion of the neuronal R_in_ and TAU by simple depolarization

R_in_ changes with voltage with current injection (Fig. 8*A*), in a voltage range from −90 mV to −15 mV. R_in_ is reduced on both ends of this voltage range (Fig. 8*A2*), likely due to activation of voltage-gated conductances (K^+^, Ca^2+^ and HCN channels; but not Na^+^ channels -- experiments done in TTX). Since R_in_ alters TAU (TAU = Rm x Cm), TAU will also change across this voltage range, as we demonstrated experimentally (Fig. 8*A3*).

### Membrane time constant TAU

The membrane charging curve was fitted with an exponential function, and the reported TAU values should be interpreted as “apparent TAU in the presence of voltage-dependent conductances”, as discussed by (Koch et al., 1996).

Strong activation of spatially distributed synaptic inputs transiently increases neuronal membrane conductance, thus lowering TAU (Bernander et al., 1991). Here we demonstrated similar effects of clustered synaptic inputs that produced dendritic plateau potential: TAU was shortened during the plateau (Fig. 6*A* model; Fig. 6*B* experiment). During the plateau potential, TAU was affected by both glutamate-induced decrease in R_in_, and to a lesser extent, by activation of voltage gated conductances. These two factors force the neuronal charging curve to reach its steady state sooner (shorter TAU) during the plateau compared to before the plateau (Fig. 6, *B2, C2* and D2).

The impact of membrane voltage alone on R_in_ or TAU was addressed in experiments. When pyramidal neurons were depolarized by dendritic plateau potentials, the changes in R_in_ and TAU were more pronounced then when the same neurons were depolarized by simple current injections (Fig. 8, compare *A* vs *B*).

### Synaptic responses are more effective during plateau potential

The most obvious factor increasing spike probability (in response to EPSPs) is the depolarization itself, which shifts the somatic (and axon initial segment) potential ∼20 mV closer to spike threshold (Fig. 3*C1*). An additional factor increasing spike probability is the increased speed of response due to decreased TAU in the cell body (Fig. 6). EPSPs with faster rise times are more powerful activators of the voltage-gated sodium current due to the sodium channel activation kinetics being faster than inactivation kinetics (Hodgkin and Huxley, 1990; Wickens and Wilson, 1998; Azouz and Gray, 2000). Shortening of TAU facilitates EPSP-induced sodium channel activation, reducing delay to threshold. However, at the same time, shortening of TAU narrows the integration time window and reduces EPSP summation. These are two disparate faces of the same biological process.

The impact of the glutamate-mediated dendritic plateau potential on the incoming EPSPs was experimentally studied in thalamocortical (TC) neurons of the dorsal lateral geniculate nucleus (Augustinaite et al., 2014). Dendritic plateau potentials evoked in distal dendritic segments of TC neurons enhanced the retinocortical transmission (proximal dendrite) by lifting the cell body membrane potential toward the threshold for action potential generation, and thereby increasing the probability for spike generation by the synaptic input from the retinal afferents (Augustinaite et al., 2014). During visual perception tasks, the feedback corticothalamic inputs arriving at distal dendritic segments of TC neurons can use dendritic plateau potentials for controlling the efficacy of retinal inputs arriving at proximal dendritic segments of TC neurons. A burst of corticothalamic inputs, which does not evoke dendritic plateau potential, but rather a strong synaptic potential, would drastically distort the relative amplitudes and relative timings inside the train of retinal inputs. The steady state phase of the plateau potential (plateau phase), on the other hand, provides a clean change in the input gain, without affecting the relative amplitudes or relative timing of the individual retinothalamic inputs.

### Timing of individual inputs

During the plateau, relative timing of sequential inputs may be preserved, as suggested by Augustinate et al. (2014). However, we found dramatic shortening of EPSP-to-AP delay during plateau potentials (Fig. 9*F*). Our current modeling data indicate that during plateau potential, a cortical pyramidal neuron can follow significantly faster changes (or oscillations) occurring in afferent neural networks, as can the very same neuron before the onset of the dendritic plateau potential.

Interestingly, dependence of dT on the Vm-t was very similar between “induced voltage change” and “plateau” (Fig. 9*H*) despite the fact that TAU differed across these 2 conditions. dT (time from input to AP) was only weakly influenced by TAU due to the countervailing effects mentioned above. The rise time of an EPSP is limited by the dynamics of the synaptic current (e.g. rise time of the current waveform) and by the membrane capacitance (Cm), but not by TAU. In pyramidal cells receiving large number of ESPS on proximal dendrites, the time delay between onset of EPSP barrage and AP generation (dT) is notably shorter than the TAU (Koch et al., 1996). Depending on the synapse clustering in space (segregation) and in time (synchronization), the threshold for spike generation can be reached in a fraction of TAU. For example, in a neuron whose TAU was found to be 20 ms, a strong and synchronous synaptic input can generate an AP only 3 ms after the onset of the EPSP barrage. On the other hand, if the EPSP input is temporally and/or spatially becoming more and more dispersed (in particular toward distal synapses) then dT will begin to be more and more influenced by the TAU (Koch et al., 1996).

### Limitations of the model

Our model is currently a special purpose model with a focus on the phenomenology of dendritic plateaus in the context of basilar and oblique dendrites. The model has been tuned to accurately match these, and to also match prior data on back-propagating APs (bAPs) in basilar dendrites, with and without channel blockers. Further extension and tuning of the model will be needed to also reflect bAPs, and synaptic integration at the apical nexus and tufts above nexus (Short et al., 2017). Additional improvements will allow us to combine this model with our prior models of corticospinal (thick-tufted) and corticostriatal (thin-tufted) Layer 5 pyramidal cells (Neymotin et al., 2017; Dura-Bernal et al., 2018).

### Dendritic plateau potentials in vivo

One major concern with experiments performed in brain slices is that dendritic signals observed ex vivo may not exist in living animals. Here, however several studies support the existence of dendritic plateau potentials in vivo (Lavzin et al., 2012; Xu et al., 2012; Smith et al., 2013; Gambino et al., 2014; Cichon and Gan, 2015; Du et al., 2017; Ranganathan et al., 2018). Significantly, two recent studies (Moore et al., 2017; Kerlin et al., 2019) reported in vivo plateau potentials with characteristics directly comparable to the biophysics of the plateau potentials discussed in the present study. Glia-encapsulated-tetrode in vivo recordings showed large-amplitude, sustained plateau depolarizations (lasting several hundred milliseconds) accompanied by sodium spikes (ref. (Moore et al., 2017), their figure 4). These dendritic plateau depolarizations are directly comparable to the data we show in Figs 1 – 5. However, in the current study, we find that only the first sodium spikelet originates in distal dendrite (Fig. 1, *C* and *D*), whereas the in vivo results showed distal-dendrite origin for the majority of the recorded fast spikelets.

High-resolution 3D calcium imaging of pyramidal neuron dendrites, out to 300 μm from the cell body, have been performed in motor cortex during a tactile decision-making task (Kerlin et al., 2019). This study demonstrated branch-specific dendritic events similar in dynamics, duration (∼1,000 – 2,000 ms) and spatial spread to NMDA-mediated plateaus described previously using calcium imaging in brain slices (Milojkovic et al., 2007; Major et al., 2008; Augustinaite et al., 2014). The longer duration reflects the fact that dendritic calcium waveform is 3 - 6 times longer than the voltage plateau when measured with voltage and calcium imaging in the same dendrite (Milojkovic et al., 2007). Such prolonged depolarization and calcium rise can contribute to synaptic plasticity (Takahashi and Magee, 2009; Brandalise et al., 2016). The authors of the in vivo study argued that long-lasting dendritic excitation could be part of the cellular mechanism of short-term memory (Kerlin et al., 2019). Short-term electrical activity is perhaps useful for storing sensory experiences from the recent past to guide present decisions and actions (Leavitt et al., 2017). Sustained depolarizations in response to strong glutamatergic input (Larkum et al., 2009; Kerlin et al., 2019), an inherent property of basal, oblique and apical tuft branches, will keep the neuron in a brief spiking mode (brief persistent activity), which is thought to be a correlate of a short-term (1 - 2 sec) dynamic memory (Goldman-Rakic, 1995; Leavitt et al., 2017). This view is in line with the idea that dendritic plateau potentials change the state of neurons for hundreds of milliseconds, by: [i] bringing their membrane potential closer to AP firing threshold (Fig. 5), [ii] enhancing the efficacy of synaptic integration (Fig. 9B), [iii] shortening the EPSP-to-AP time (Fig. 9F), and [iv] improved detection of faster network rhythms (Fig. 9, C and D).

## Acknowledgements

We are grateful to Corey Acker and Guy Major for providing their model code, to Chun Bleau (Redshirt Imaging) and Charlie Bleau (SciMeasure Analytical Systems) for assistance with imaging equipment. This work was supported by National Institute of Mental Health, MH063503, MH109091 and MH086638; National Institute of Neural Disorders and Stroke, NS11613; National Institute of Biomedical Imaging and Bioengineering, EB022903 and EB017695.

## Competing Interests

The authors declare no competing financial interests.

## Supplemental Data

https://figshare.com/articles/Suppl_Figs_v02_pdf/11301020

https://doi.org/10.6084/m9.figshare.11301020.v1

Supplemental data is comprised of seven figures with legends, and deposited at figshare.com (file name: Suppl Figs_v02.pdf). Both links (above) point to the same file.

## Notes

https://figshare.com/articles/Suppl_Figs_v02_pdf/11301020

### Competing Interest Statement

The authors have declared no competing interest.

### Summary of Updates

New experimental data is included. Model predictions were tested in real neurons.

